# Conserved cerebellar rhombic lip compartmentalization and *Eomes* regulatory networks govern unipolar brush cell development

**DOI:** 10.64898/2026.07.27.740783

**Authors:** Vuslat Akçay, Lisa M. Spänig, Wouter-Michiel A.M. Vierdag, Piyush Joshi, Patricia Benites Gonçalves da Silva, Angelina Sanderson, Robert Reinhardt, Laura Sieber, Nina Hofmann, Jana Nolle, Franziska Schelb, Marc Zuckermann, Henrik Kaessmann, Stefan M. Pfister, Adriana Sánchez-Danés, Mari Sepp, Sinem K. Saka, Lena M. Kutscher

## Abstract

The rhombic lip (RL) gives rise to all cerebellar glutamatergic cell types, including unipolar brush cells (UBCs). Disruptions to UBC development can lead to the neurodevelopmental disorder Dandy-Walker Syndrome and the pediatric brain tumor medulloblastoma, but these diseases have not been adequately modeled in mice. To evaluate conservation of UBC development in mouse and human, we examined UBC localization, lineage decisions, and the underlying molecular mechanisms of UBC differentiation using multiplex immunofluorescence and single-cell RNA-seq of wild-type and conditional knockout animals of the primary UBC transcription factor *Eomes*. Similar to the human RL, the murine RL is molecularly compartmentalized, cycling EOMES+ UBC progenitors are highly abundant, and persist after birth. *Eomes* regulates the transcriptional networks important for UBC differentiation and migration, but not UBC fate. Overall, our findings suggest that murine UBC development recapitulates many features of human UBC development, with EOMES playing a central role in UBC maturation.

## INTRODUCTION

The cerebellum is the brain region responsible for balance and coordination, and it has recently been linked to higher cognitive functions, including language, verbal and spatial memory, and executive functioning^1^. Its protracted development makes it highly susceptible to neurodevelopmental disorders and disease^2^. In particular, the rhombic lip (RL) is the site of origin for the neurodevelopmental disorder Dandy-Walker Syndrome and the pediatric brain cancer medulloblastoma, specifically subgroups 3/4^3–5^. Notably, neither Dandy-Walker Syndrome nor Group 4 medulloblastoma have been faithfully modeled in animal models, suggesting that some aspects of RL development may be unique to humans.

The human RL has a proposed unique morphological compartmentalized structure, split into a ventricular zone (RL-VZ), characterized by expression of the transcription factors (TFs) *LMX1A* and *SOX2*, and a subventricular zone (RL-SVZ) comprised primarily of cycling UBC progenitors, characterized by *LMX1A* and *EOMES* expression, physically separated by a vascular bed^4,6^. The murine RL is also molecularly compartmentalized at embryonic (E) day 15, divided into inner and outer compartments separated by an EOMES+ UBC zone^7^. Because murine compartmentalization has only been examined at E13, E15, and E18^7^, it is unclear for how long this structure persists and which cellular features are shared with the human RL. Furthermore, whether a cycling EOMES+ UBC progenitor exists in mice and its relative abundance are debated^8,9^, a question relevant to whether diseases arising from such progenitors (*e*.*g*., Group 3/4 medulloblastoma) can even be modeled in mice. Finally, while *Eomes* is required for proper differentiation of murine UBCs^10^, the molecular mechanisms behind this process are unknown.

To resolve these questions, we characterized the murine RL across development using multiplex immunofluorescence (Immuno-SABER^11^) and single-cell RNA-sequencing, comparing with human references where possible. We found that the murine RL is compartmentalized over two prenatal days, comprised primarily of cycling UBC progenitors, and that some of these progenitors persist even after birth. We also investigated the molecular role of *Eomes* in UBC lineage development using conditional knockout (cKO) mouse models. Through the integration of developmental trajectory and gene regulatory network analyses, we defined the cellular lineage and developmental progression of murine UBCs and determined how *Eomes* contributes to the transcriptional programs underlying UBC specification and differentiation, specifically by maintaining *Lmx1a* transcriptional networks, the main UBC cell fate determining TF. Overall, our data reveals a higher degree of similarity between mouse and human UBC development than has previously been recognized, suggesting that the failure to model specific cerebellar diseases in mice is unlikely due to differences in cell types or structures, but may instead reflect other human-specific features, such as genome architecture or prolonged gestation.

## RESULTS

### Molecular compartmentalization of the murine and human RL is conserved

To examine RL compartmentalization, we compared mouse and human samples using Immuno-SABER (Figure S1A). In the developing mouse RL, we analyzed five timepoints: E13, E15, E16, E17, and P0 (Figure 1A). We visualized seven key developmental TFs (EOMES, LMX1A, BARHL1, ATOH1, PAX6, OTX2 and SOX2), three UBC receptor markers (mGluR1α, Calretinin/CR, and TRPC3), and the proliferation marker KI67 (Figure S1A; Table S1; see *Methods*). Due to antibody limitations, we were unable to stain for ATOH1 and LMX1A at all timepoints.

**Figure 1:**
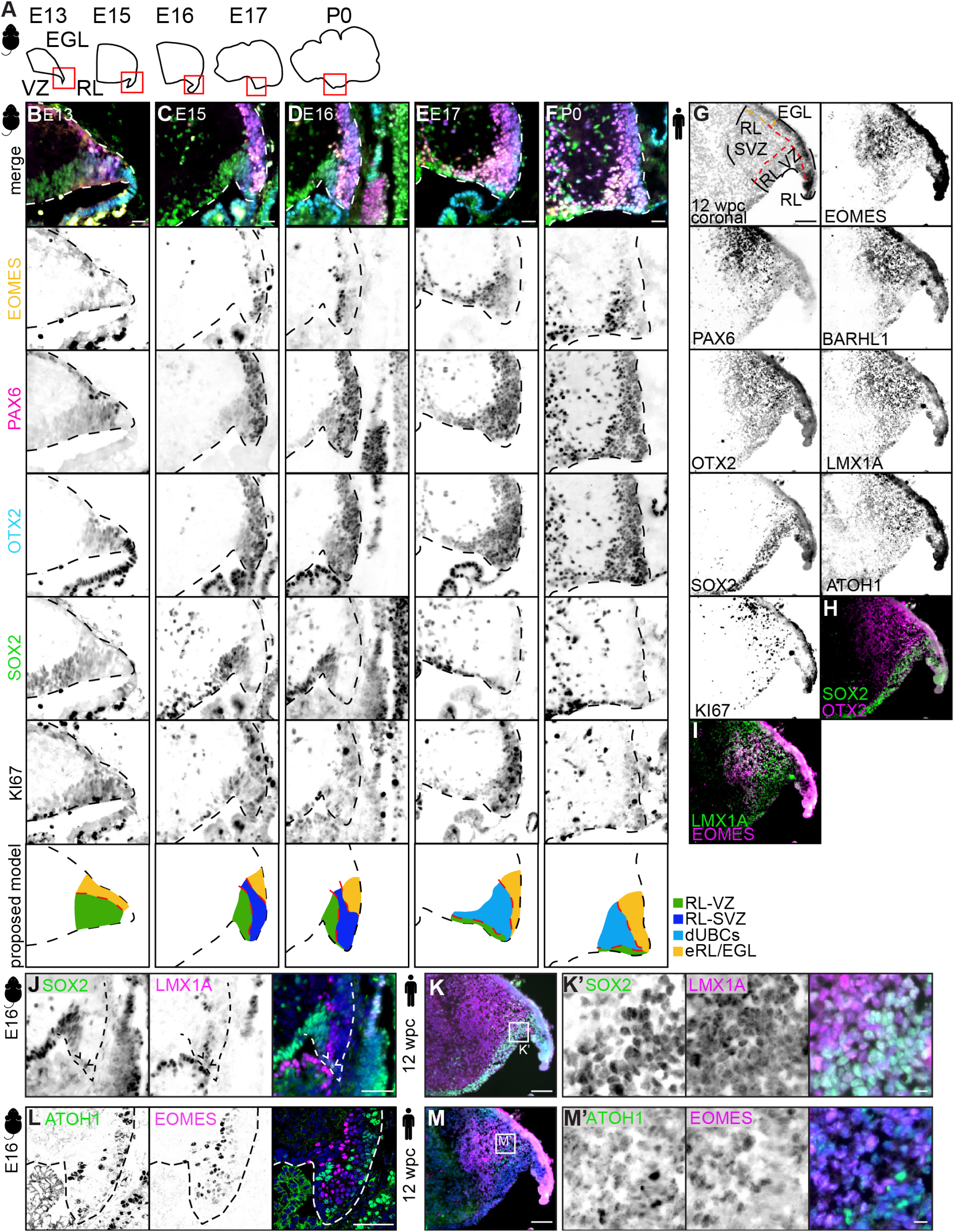
Developing mouse RL is molecularly compartmentalized. **A**) Overview of regions imaged using Immuno-SABER. **B-F**) Immuno-SABER of EOMES (yellow), PAX6 (magenta), OTX2 (cyan), SOX2 (green), and Ki67 (gray) at B) E13, C) E15, D) E16, E) E17, and F) P0. Dotted line, cerebellar anlage. Green, RL ventricular zone (RL-VZ). Dark blue, RL sub-ventricular zone (RL-SVZ). Light blue, differentiating UBCs (dUBCs). Yellow, external RL (eRL) and external granule layer (EGL). Scale bar, 25µm. **G**) Immuno-SABER of 12 weeks post-conception (wpc) human sample with respective antibodies. Scale bar, 100µm. **H**,**I**) Overlays of H) SOX2 (green) and OTX2 (magenta), I) EOMES (magenta) and LMX1A (green). **J**) Murine E16 RL immunofluorescence (IF) of SOX2 (green) and LMX1A (magenta). Scale bar, 50µm. **K**) Immuno-SABER overlay of SOX2 (green) and LMX1A (magenta) from 12 wpc sample. Scale bar, 100µm. K’, zoom of box in K. Scale bar, 10µm. **L**) Murine E16 RL IF of ATOH1 (green) and EOMES (magenta). Scale bar, 50µm. **M**) Immuno-SABER overlay of ATOH1 (green) and EOMES (magenta) from 12 wpc sample. Scale bar, 100µm. M’, zoom of box in M. Scale bar, 10µm.

We first examined the spatial localization of SOX2 (VZ marker), PAX6 (glutamatergic marker), OTX2 and EOMES (granule cell (GC) and UBC lineage markers), and KI67 (Figure 1B-F). While both the RL-VZ and RL-SVZ compartments comprised PAX6+, OTX2+, and KI67+ cells, SOX2 and EOMES expression separated these niches, indicating clear molecular compartmentalization of the murine RL (Figure 1B-F). This compartmentalization was most prominent at E15 and E16.

We next performed Immuno-SABER on a fetal human cerebellar section at 12 weeks post-conception (wpc; Figure 1G), corresponding to ∼E16 in mice^12^. We identified a SOX2+LMX1a^lo^ RL-VZ (Figure 1G,H) and an EOMES+LMX1A+ RL-SVZ (Figure 1G,I), consistent with expected molecular compartmentalization^6^. At E16 in mice, several SOX2+LMX1A+ double-positive cells were detected (Figure 1J), but these cells comprised the entire RL-VZ in the human sample, as expected^6^ (Figure 1K,K’). Although previous studies have shown that a vascular bed separates these niches^6,13^, it was not structurally apparent in our sample. We examined expression patterns of the erythroid and endothelial genes *HBE1* and *KDR* derived from published spatial transcriptomic data^12^ from an adjacent section of the same human sample to confirm this finding (Figure S1B,B’). Similar to the mouse cerebellum at E15 (Figure S1C), blood vessels were not enriched in the RL, and their density was similar across the cerebellum in both species (Figure S1B,C). We note that our human section is coronal, whereas previous analyses used sagittal sections^4,13^, which may account for these differences. Overall, these data suggest that while both human and murine RL are molecularly compartmentalized into RL-VZ and RL-SVZ, with similar expression patterns of core TFs in both species, neither species displayed morphological compartmentalization in the form of a vascular bed, at least in our samples.

Next, we analyzed whether GC_UBC progenitors, defined by large-scale single-cell atlasing studies^12^ as co-expressing the glutamatergic lineage marker *Atoh1* and the UBC marker *Eomes*, also co-express these TFs at the protein level, or whether their co-occurrence in transcriptomic data reflects transcriptional priming toward discrete cell fates^14^. At our earliest timepoint, E13 in mice, the glutamatergic lineage TF ATOH1 was expressed primarily in the growing external granule layer (EGL; Figure S1D,E), and the majority of EOMES+ cells resided in the nuclear transitory zone (NTZ), with minimal overlap with LMX1A+ cells (Figure S1F). At E13, several EOMES+ATOH1+ cells were observed in the RL (Figure S1E), while at E16 these proteins did not overlap (Figure 1L). In the human sample, co-expressing cells were abundant (Figure 1M,M’). These data demonstrate that GC_UBC progenitors indeed co-express these TFs at the protein level in both species, and they are more abundant in humans.

To corroborate these findings, we reanalyzed published^12^ cerebellar murine and human single-cell sequencing data to determine the relative abundance of GC_UBC progenitors and differentiating UBCs. At 11 wpc/E15, we again observed these cells in both species, with a relatively greater abundance within the GC and UBC lineage cells in humans compared with mice (Figure 2A,B), suggesting the UBC lineage is expanded during human RL development. We propose that the relative abundance of these cells is the key distinction between murine and human development.

**Figure 2:**
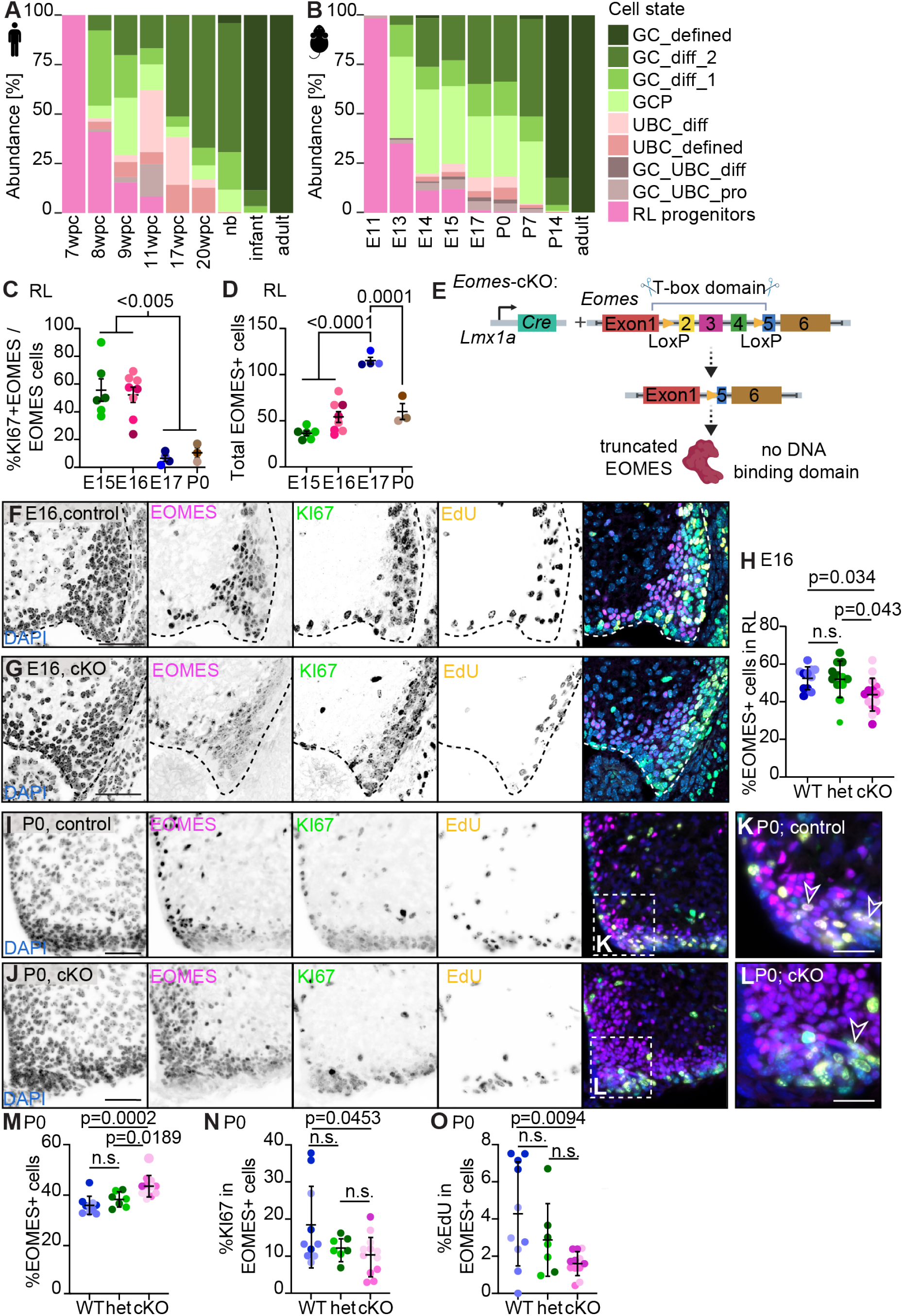
Cycling EOMES+ cells are highly abundant E16, mimicking the RL-SVZ in humans. **A**,**B**) Percent abundances of each cell state derived from published single-cell developing RL lineage data^12^. Corresponding developmental stages matched between A) human and B) mice. wpc, weeks post-conception. nb, newborn. E, embryonic day. P, postnatal day. Sampling after 11 wpc comes from tissue fragments and may not be fully representative. GC, granule cell. Diff, differentiating. GCP, granule cell progenitor. UBC, unipolar brush cell. GC_UBC_pro, granule cell and unipolar brush cell progenitor. RL, rhombic lip. **C**) Quantification of Immuno-SABER %KI67+EOMES/EOMES+ RL cells over time. Dots colored by developmental time point. Ordinary one-way ANOVA. Multiple comparisons revealed E15 and E16 as significantly different from E17 and P0. **D**) Quantification of Immuno-SABER total EOMES+ RL cells over time. Dots colored by developmental time point. Ordinary one-way ANOVA. Multiple comparisons revealed E15, E16, and P0 as significantly different from P0. **E**) Overview of cKO mouse line. **F**,**G**) Representative IF of E16 RL of F) control and G) cKO animals. DAPI, blue. EOMES, magenta. KI67, green. EdU, yellow. Scale bar, 50µm. **H**) Quantification of E16 %EOMES RL cells. Dots of same shade were quantified from different cerebellar sections of same mouse. N=3-4 sections from 3 animals. Ordinary one-way ANOVA. n.s., non-significant. WT, wild-type. het, heterozygous loss of *Eomes*. cKO, conditional knockout. **I**,**J**) Representative IF of P0 cerebellum of I) control and J) cKO animals. DAPI, blue. EOMES, magenta. KI67, green. EdU, yellow. Scale bar, 50µm. **K**,**L**) Zoom image from I and J, respectively. Carets represent EOMES+EdU+ cells. Scale bar, 25µm. **M-O**) Quantification of M) %EOMES cells, N) proliferating %KI67+EOMES/total EOMES cells, O) %EdU+EOMES+/total EOMES cells in 200×200µm area around former RL at P0. Only DAPI+ cells within the cerebellum were counted. Ordinary one-way ANOVA. n.s., non-significant. Dots of same shade were quantified from different cerebellar planes of same mouse. N=3-4 sections from 3 animals.

Finally, using our Immuno-SABER data, we characterized migration and differentiation patterns of EOMES+ cells in mice. By E16 and E17, EOMES+LMX1A+ UBCs were abundant in the RL (Figure S1G,H), and the UBC receptors CR and TRPC3 were already detectable at E17 (Figure S1I,J), with expression increasing at P0 (Figure S1K,L). At E17, we observed two streams of EOMES+ cells emerging from the RL: an EOMES^hi^LMX1A+ population migrating through the developing white matter, and an EOMES^lo^LMX1A-population along the EGL, (Figure S1G). At P0, this EOMES^lo^ population was present along the inner EGL (iEGL) of the posterior cerebellum, co-expressing PAX6 and BARHL1 (Figure S2A,B). As these cells lacked LMX1A, we hypothesize that they are migrating EOMES+ GCs^15^.

Together, these findings suggest that the overall molecular structure and cellular composition of the murine and human RL are conserved, with the greatest similarity in compartmentalization occurring around E16.

### Cycling EOMES+ progenitors peak at E16

It has been suggested that cycling EOMES+ RL progenitors are rare in mice^4,5^: some reports question their existence entirely^8^, while others demonstrate that cycling *Eomes*+/EOMES+ cells are present at E16^9,16^. These studies, however, only examined single timepoints or did not characterize UBC development beyond E16. Therefore, we quantified EOMES+KI67+ co-expression in the RL and cerebellum from E15 to P0. Cycling EOMES+ cells represented approximately 50% of the total EOMES+ RL population (Figure 2C) and ∼20% in the cerebellar anlage (Figure S2C) at both E15 and E16. By E17 and P0, this fraction fell to ∼5% in the RL (Figure 2C). As molecular compartmentalization persisted at E17, but most cells no longer expressed KI67, we designated this region as differentiating UBCs (dUBCs), rather than RL-SVZ (Figure 1F). The total EOMES+ RL population, including both cycling and non-cycling cells, increased steadily from E15 to E17 before declining at P0, as cells migrated away (Figure 2D;S2D). Together, these data confirm that E16 represents the peak of cycling EOMES+ progenitor abundance, with some UBC progenitors persisting beyond birth.

### Loss of Eomes in the Lmx1a-Cre lineage does not affect UBC progenitor specification

To investigate the functional role of *Eomes* in UBC progenitor fate specification, we generated an *Eomes* cKO animal line. We crossed a floxed *Eomes* allele, which creates an in-frame deletion lacking most of the DNA binding domain^17^, with *Lmx1a-Cre*^18^ animals (Figure 2E). *Lmx1a-Cre* is expressed in the posterior cerebellum^18^, where UBCs are localized. This cross enables visualization of both the wild-type protein and the truncated mutant protein using a C-terminal-targeting antibody, allowing us to visualize mutant EOMES protein expression in *Eomes*^*cKO*^ genotyped mice. Mutant protein was detected at E16 and P0 (Figure 2F-J), but EOMES+ signal was not present at P28 in *Eomes*^*cKO*^ mice (Figure S2E,F) and mature UBCs were ablated (Figure S2G,H). Overall cerebellar structure and size were comparable at P28 (Figure S2I-K).

At E16, quantification of EOMES+ cells in the RL showed that these cells were reduced in the *Eomes*^*cKO*^ background (Figure 2H; p=0.034, t-test); however, positive staining of mutant EOMES indicated that UBC progenitors were properly specified in *Eomes*^*cKO*^ genotyped mice. To assess whether EOMES+ mutant UBC progenitors were still mitotically active, we injected the thymidine analog EdU into pregnant dams at E16 and harvested brains 2h later. Quantification of KI67+EdU+EOMES+ cells in the RL revealed no differences in the proportion of cycling UBC progenitors (KI67+EOMES+; Figure S2L) or EdU incorporation (∼20% of KI67+EdU+EOMES+ cells in active S-phase at E16; Figure S2M). Total DAPI+ cell number in the RL at E16 was unchanged (Figure S2N).

We performed a similar set of experiments at P0 (Figure 2I,J). EOMES+ cells in active S-phase were found in both *Eomes*^*WT*^ and *Eomes*^*cKO*^ animals at P0 (Figure 2K,L). Notably, we observed an increase in the proportion of EOMES+ cells at the former RL in *Eomes*^*cKO*^ animals (Figure 2M; p=0.0002, t-test), alongside a modest decrease in the proportion of cycling UBC progenitors (Figure 2N; p=0.0453, t-test). Approximately 3-4% of EOMES+ cells were in active S-phase (EdU+) in controls, compared with <2% in *Eomes*^*cKO*^ animals (Figure 2O; p=0.0094, t-test). Together, these data demonstrate that actively dividing EOMES+ progenitors are present at both E16 and P0, and that the loss of *Eomes* DNA-binding activity may affect the abundance and/or localization of UBCs in this lineage.

### Eomes+ UBCs and a subset of Eomes+ GCs derive from an Lmx1a-expressing progenitor

To better understand the lineage relationships between EOMES+ UBCs and EOMES+ GCs, we performed single-cell RNA-sequencing on enriched GC_UBC lineage cells in three separate mouse models (Figure 3A). The first mouse strain, *Eomes-IRES-GFP* knock-in reporter line^19^, was used to provide data on *Eomes+* cells in wild-type animals at baseline. In this line, GFP is expressed from the *Eomes* locus without disrupting the endogenous gene. GFP signal overlapped with EOMES antibody staining at E16 (Figure S3A,B) and P0 (Figure S3C,D), indicating that EOMES expression could be followed with the GFP signal.

**Figure 3:**
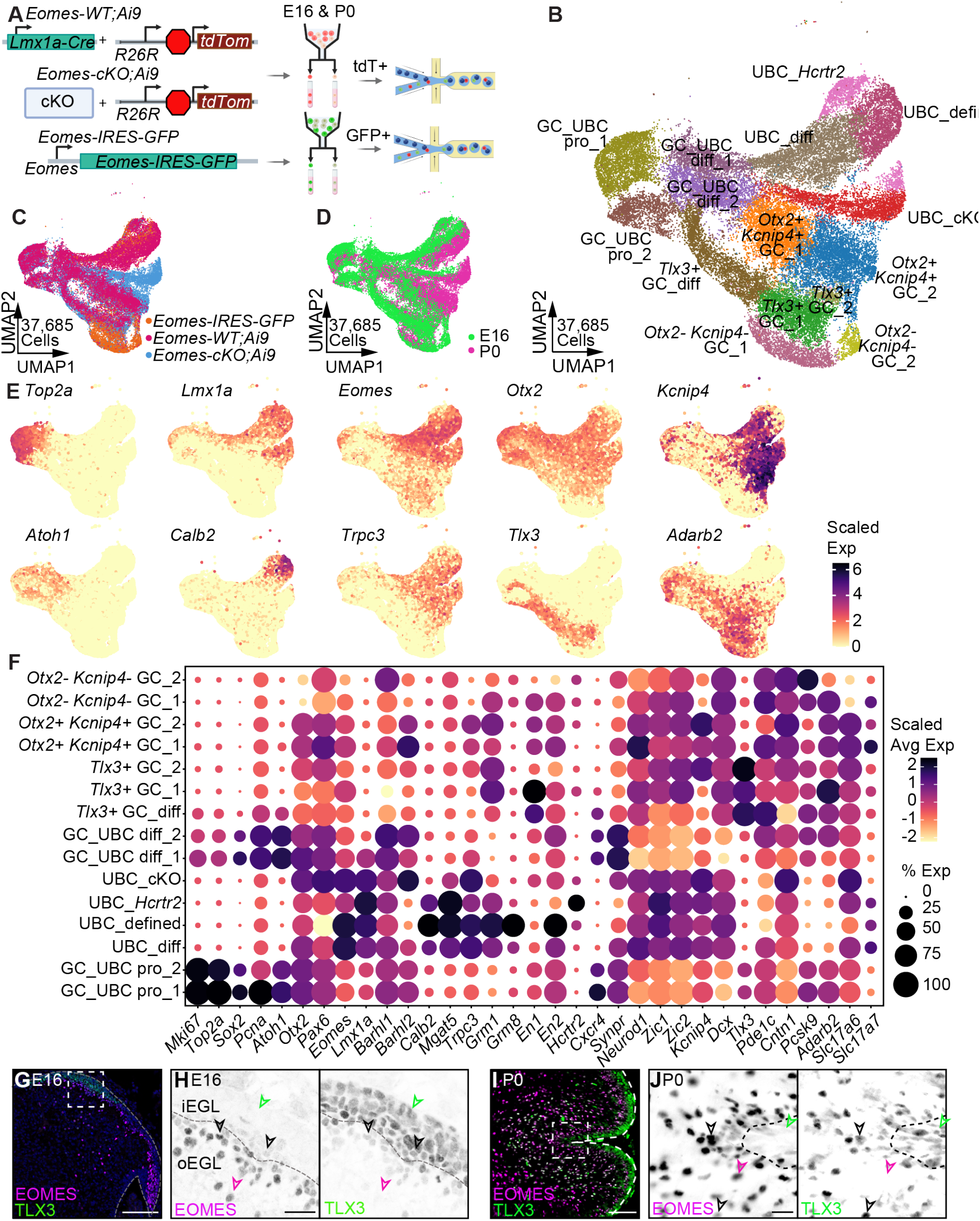
EOMES+ UBCs and a subset of EOMES+ GCs are derived from an *Lmx1a*-expressing progenitor. **A)** Overview of single-cell RNA-sequencing experiment. **B-D**) Single-cell RNA-seq samples of sorted GFP+ cells from *Eomes-IRES-GFP* cerebellum and sorted tdTomato+ cells from *Eomes-WT;Ai9* and *Eomes-cKO;Ai9* cerebellum, at E16 and P0, filtered for the GC_UBC lineage and colored by B) cell type and states, C) sample origin, and D) age. **E**) Scaled expression of cell state and TF marker genes, visualized by feature plots. **F**) Scaled marker gene expression of cell types and states, visualized by dot plot. **G**) Representative IF image of wild-type E16 cerebellum. DAPI, blue. EOMES, magenta, TLX3, green. Scale bar, 100µm. **H**) Zoomed in image of boxed area in G. Scale bar, 20µm. iEGL, inner external granule layer. oEGL, outer external granule layer. **I**) Representative IF image of wild-type P0 cerebellum. EOMES, magenta, TLX3, green. Scale bar, 100µm. **J**) Zoomed in image of boxed area in I. Scale bar, 20µm. Black arrow, co-expressing cells. Pink arrow, EOMES-expressing cell. Green arrow, TLX3-expressing cell.

In two additional mouse strains, we crossed *Lmx1a-Cre* animals (± *Eomes*^*floxed*^ allele) to a *LoxP-STOP-LoxP-TdTomato* reporter cassette (*Ai9*), labeling all cells derived from the *Lmx1a-Cre* lineage. GFP+ (*Eomes-IRES-GFP*) or TdTomato+ (*Eomes-WT;Ai9* and *Eomes-cKO;Ai9*) cells were sorted from dissected cerebella at E16 and P0 for single-cell RNA-seq, resulting in six samples in total. From the single-cell data, we identified 19 clusters belonging to 15 distinct GC_UBC lineage cell types and states (Figure 3B), after removal of non-GC_UBC lineage cells (Figure S3E,F). Within the control lineages, both *Eomes-IRES-GFP* and *Eomes-WT;Ai9* cells contributed to nearly all GC_UBC clusters at both E16 and P0 (Figure 3C,D;S3G,H).

To assign cluster identities on the composite data set, we examined known marker genes for cell state (*e*.*g. Top2a, Mki67*), cell fate (*e*.*g. Atoh1, Eomes, Lmx1a*) and receptor markers (*e*.*g. Calb2, Trpc3*)^12^ (Figure 3E;S3I). We identified progenitor clusters (GC_UBC pro and GC_UBC diff), UBC lineage clusters (UBC_diff, UBC_defined, UBC_*Hcrtr2*) and GC lineage clusters (*Tlx3+* GCs, *Otx2+Kcnip4+* GCs, *Otx2-Kcnip4-* GCs; Figure 3F). The *Otx2-Kcnip4-* GCs cluster almost entirely comprised *Eomes-IRES-GFP* cells (Figure S3J), suggesting that *Otx2-Kcnip4-* GCs do not derive from an *Lmx1a+* progenitor. Conversely, both *Tlx3+* GCs and *Otx2+Kcnip4+* GCs were found in both control samples, suggesting that these *Eomes+* GC subtypes arise from an *Lmx1a-*expressing progenitor. Within the UBC lineage, differentiating and defined UBCs expressed receptor genes characteristic of both UBC subtypes (Figure 3E;S3J) and were found in both control samples.

As *Eomes*+ GCs have only recently been described^15^, we validated the presence of TLX3+EOMES+ co-expressing cells in the wild-type developing cerebellum by immunofluorescence. EOMES+ cells were distributed across the posterior cerebellum, while TLX3+ cells were restricted to a specific region of the EGL (Figure 3G). TLX3 expression was highest in the outer EGL (oEGL), corresponding to actively cycling cells^20^, while cells co-expressing TLX3 and EOMES were largely restricted to the iEGL (Figure 3H), which is post-mitotic^20^. At P0, this pattern persisted; TLX3+ granule cell progenitors (GCPs) spanned the posterior lobe EGL (Figure 3I), while the inner granule layer (IGL) in this region contained a mix of TLX3+ only, EOMES+ only and TLX3+EOMES+ double positive cells (Figure 3J).

Overall, these data demonstrate that all *Eomes+* UBCs and a subset of *Eomes+* GCs derive from the *Lmx1a* lineage, with only *Eomes+Otx2-Kcnip4-* GCs arising outside this lineage.

### Eomes is essential for UBC maturation and migration

While examining our data, we noticed one cluster consisting primarily of mutant cells with >99% *Eomes-cKO*; *Ai9* contribution (Figure S3J). This cluster retained UBC markers including *Lmx1a* and *Eomes* (Figure 3F), as well as markers of differentiating GC_UBC and UBC states, suggesting that these cells retain UBC lineage identity. Therefore, we called this cluster UBC_cKO.

To characterize transcriptional changes in these UBC_ cKOs compared to control UBCs, we performed differential gene expression (DEG) analysis (Figure 4A;S4A). We used an adjusted p-value <0.05 and log_2_ fold change cutoff of ±1 (Table S2). Specifically, cells from the UBC_cKO cluster and cells from UBC_diff and UBC_defined clusters were pseudobulked separately to generate control and cKO UBC populations at E16 and P0 (Figure 4A;S4A). We found that canonical UBC receptor genes, including *Calb2, Grm1*, and *Mgat5*, were downregulated in UBC_cKOs at both E16 and P0 (Figure 4A;S4A), indicating that although UBC identity was correctly specified, mutant cells failed to mature. To identify biological processes affected by *Eomes* loss, we performed Gene Set Enrichment Analysis (GSEA) and Gene Ontology (GO) analyses (Figure 4B;S4B,C; Table S3,S4). Ranking GSEA results by significance revealed that UBC_ cKO were enriched for pathways related to cell motility, locomotion, and adhesion at P0, including members of the *Spock* family (Figure 4A,B). GO analysis at P0 also revealed enrichment of signaling pathways related to UBC function and connectivity in both control and UBC_cKO clusters, including programs involved in synapse assembly and axon guidance (Figure S4B). Analysis of individual axon guidance genes showed distinct sets of enriched genes in control and UBC_cKOs, indicating extensive rewiring of axon guidance-related transcriptional programs following *Eomes* loss, including members of the *Slit* and *Semaphorin* families (Figure S4C).

**Figure 4.**
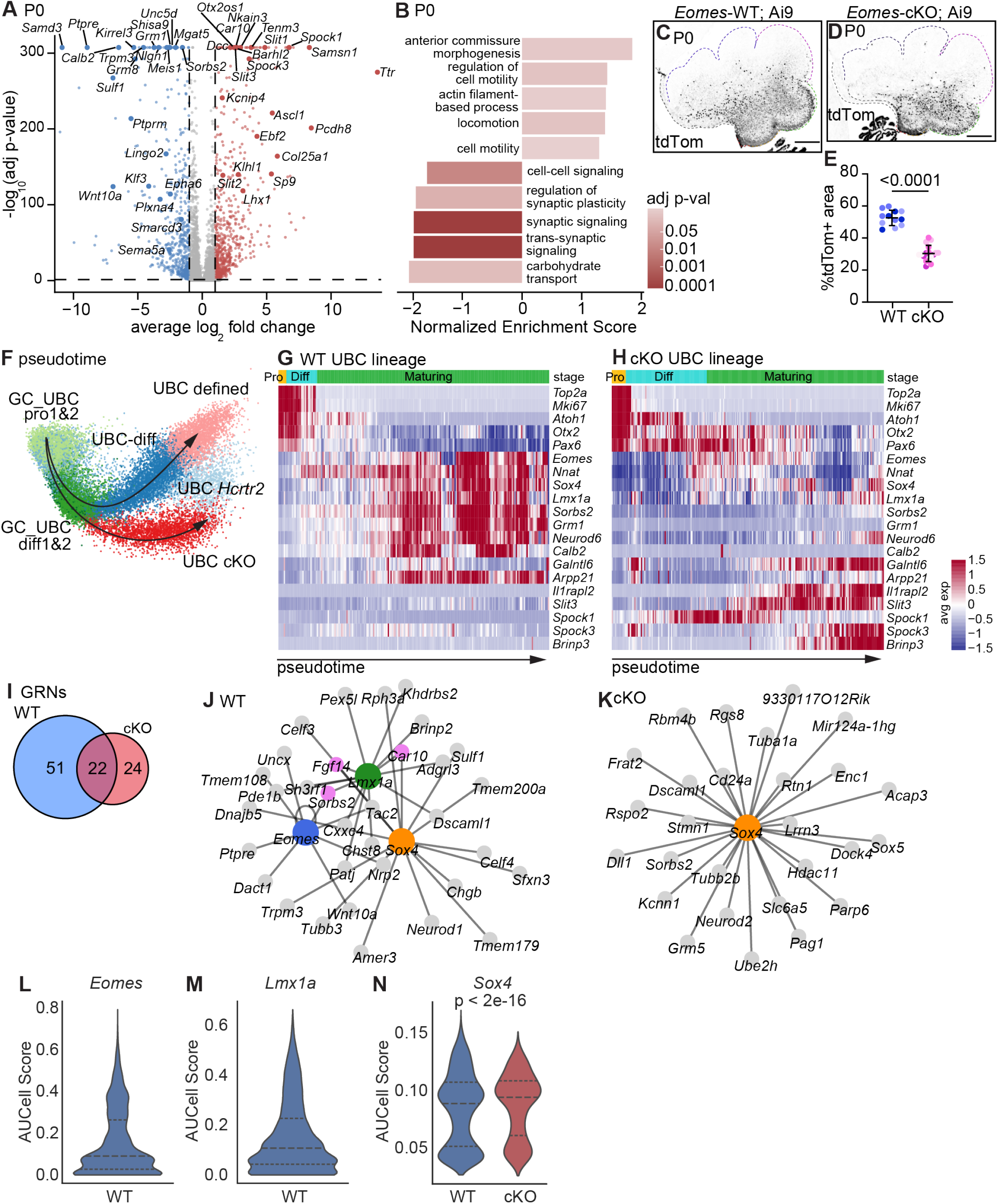
*Eomes* is required for the regulation of UBC maturation, migration and *Lmx1a* network function. **A**) Volcano plot comparing pseudobulked UBC_cKO vs UBC at P0. Blue, genes downregulated in UBC_cKO. Pink, genes upregulated in UBC_cKO. **B)** Gene set enrichment analysis showing biological processes in UBC_cKO at P0. **C**,**D**) tdTom staining in C) *Eomes-WT;Ai9* D) *Eomes-cKO;Ai9* cerebellum at P0. Scale bar, 200µm. **E**) Quantification of tdTomato+ area. T-test. Dots of same shade were quantified from different cerebellar sections of same mouse. N=3-4 sections from 3-4 animals. **F**) Pseudotime UMAP of UBC lineage using merged *Eomes-WT;Ai9* and *Eomes-cKO;Ai9* datasets. **G-H**) Expression dynamic heatmaps of early- and late-stage genes across pseudotime-ordered UBC lineage in G) *Eomes-WT;Ai9* and H) *Eomes-cKO;Ai9* cells. Pro, progenitors. Diff, differentiating. **I**) Venn diagram of unique and shared active gene regulatory networks (GRNs). **J**,**K**) SCENIC TF-target gene networks for J) *Eomes, Lmx1a*, and *Sox4*, with top predicted target genes in *Eomes-WT;Ai9* and K*) Sox4* and top predicted target genes in *Eomes-cKO;Ai9* cells. Grey nodes, predicted target genes. Edges, predicted regulatory interactions. **L)** Regulon activity score of *Eomes* in *Eomes-WT;Ai9*. Network is lost in *Eomes-cKO;Ai9* cells. **M)** Regulon activity score of *Lmx1a* in *Eomes-WT;Ai9*. Network is lost in *Eomes-cKO;Ai9* cells. **N)** Regulon activity score of *Sox4* in WT and cKO. AUCell Score, area under curve cell score.

Given the aberrant expression of migration- and axon guidance-related genes, and the increased number of EOMES+ RL cells in *Eomes*^*cKO*^ animals at P0, this accumulation likely reflects a migration defect rather than increased cell number, consistent with previous findings^10^. Using the *Ai9* reporter to trace the *Lmx1a-Cre* lineage at P0 (Figure 4C,D), we found that tdTomato+ cells occupied a larger area in control animals than in *Eomes*^*cKO*^ animals (Figure 4E), suggesting that *Eomes*^*cKO*^ RL cells fail to migrate efficiently. Together, these results demonstrate that *Eomes* loss disrupts axon guidance and migration gene expression, preventing UBC maturation and proper migration.

To further assess the developmental state of UBC_cKOs, we performed trajectory inference and pseudotime analysis on the combined *Eomes*^WT^ and *Eomes*^cKO^ *Ai9* datasets. The resulting trajectory showed that lineage divergence following *Eomes* loss occurs after the differentiating GC_ UBC state, with cells branching towards either maturing wild-type UBCs or UBC_cKOs (Figure 4F). The shared UBC_ *Hcrtr2* state occupied an intermediate position between these trajectories, suggesting that *Eomes* loss may only partially affect differentiation of this *Eomes*^*lo*^*-* expressing population^12^. We next ordered the UBC lineage by pseudotime and examined stage-specific marker genes (Figure 4G,H). During normal maturation, UBCs transition from expressing early developmental markers (*Atoh1, Otx2, Pax6*) to expressing more specialized markers (*Eomes, Nnat, Sox4, Lmx1a, Sorbs2*, and *Neurod6*; Figure 4G). *Eomes*^cKO^ cells, however, failed to downregulate the early TFs *Otx2* and *Pax6* (Figure 4H), possibly maintaining UBC_cKOs in a less differentiated state. Maturing UBC_cKOs also largely lacked mature UBC markers and instead aberrantly upregulated ECM remodeling genes (Figure 4G,H). Overall, pseudotime analysis indicates that the UBC_cKO lineage stalls at an earlier, aberrant developmental stage compared with control, allowing misexpression of genes not typically present within the UBC lineage.

### Eomes is required for the maintenance of Lmx1a-associated gene regulatory networks

To investigate the regulatory mechanisms underlying these defects, we performed TF-gene network analysis using Single-Cell Regulatory Network Inference and Clustering (SCENIC)^21^ on *Eomes-*WT;*Ai9* and *Eomes-*cKO;*Ai9* cells. SCENIC analysis identified 73 active gene-regulatory networks (GRNs) in WT and 46 active GRNs in cKO cells, of which 22 were shared, indicating a substantial reduction in the regulatory network landscape following *Eomes* loss (Figure 4I).

We next examined individual GRN activities (called regulon activities) in greater detail. As expected, the *Eomes* regulon was absent in cKO cells (Figure 4J-L;S4D). Activity of the *Lmx1a* regulon was also lost, despite preserved *Lmx1a* transcript expression, suggesting disruption of its regulatory activity rather than basal expression (Figure 4J-K,M). Network analysis of WT cells revealed that *Eomes, Lmx1a*, and *Sox4* formed a closely connected regulatory module (Figure 4J). Although *Sox4* expression and regulon activity were retained in cKO cells, its regulatory connectivity was markedly reduced following *Eomes* loss (Figure 4K,N). We also identified *Sorbs2*, a late-stage UBC receptor gene (Figure 4G), as a predicted target of both *Eomes* and *Lmx1a* regulatory networks (Figure 4J). Together, these findings illuminate putative mechanisms of gene regulation by *Eomes* in UBC differentiation and maturation, including maintaining *Lmx1a* regulatory network activity within maturing UBCs. Furthermore, the close TF-associations between *Eomes, Lmx1a*, and *Sox4* suggest that this regulatory module may be essential to establish mature functional UBC identity.

## DISCUSSION

The RL is a key cerebellar germinal zone, with EOMES+ cells comprising an abundant cell population in both murine and human fetal development. We show that the murine and human RL share similar cell types and compartmental organization, consistent with large-scale single-cell studies demonstrating molecular similarities between cell types^6,12^. While E16 represents the peak of cycling UBC progenitor abundance in the mouse, our comparative analyses suggest that this lineage is proportionally more abundant in humans at equivalent developmental stages, especially SOX2+LMX1A+ RL-VZ progenitors and ATOH1+EOMES+ GC_UBC progenitors. We also show that cycling UBC progenitors are highly abundant in the prenatal murine RL, and also persist postnatally, indicating that even this aspect of RL development is conserved across species. Despite these similarities, other differences must exist, given that certain diseases have not yet been adequately modeled in mice. We suspect these differences are not specific to the RL itself, because although specific cell types are less abundant in mice, they are still present. We hypothesize that failure to model these diseases may instead reflect other human-specific features, such as genome architecture^22^; for example, the majority of Group 3/4 medulloblastoma tumors harbor recurrent large-scale chromosomal copy-number changes^23^ that cannot be recapitulated completely in the murine genome.

Prior work using *Nes-Cre*-mediated *Eomes* deletion demonstrated an absence of mature UBCs and impaired UBC migration in the cerebellum^10^. Our study provides new insights by dissecting the molecular basis of this defect using the more lineage-specific *Lmx1a-Cre* transgene. We find that neuronal migration pathways and axon-guidance molecules are misexpressed upon *Eomes* loss, including Slit-Robo repulsive signaling, Semaphorin signaling, and Netrin receptor expression. In the developing chick, Netrin1 promotes migration of RL-derived cells while Slit2 inhibits it^24^, suggesting that one role of EOMES may be to directly regulate these migration signaling molecules to enable maturing UBCs to exit the RL. This is consistent with the observation that UBC progenitors appear to reside in the RL for several days before initiating migration during prenatal development^16^.

Finally, our GRN analyses suggest that *Eomes* is essential to maintain an *Lmx1a*-associated regulatory network during UBC maturation. In the absence of *Eomes*, the *Lmx1a* regulon is disrupted and the *Sox4* network is altered, although both genes remain expressed. In the blastocyst, EOMES serves as a pioneering TF and chromatin remodeler that drives mesoderm and endoderm (ME) lineage specification during gastrulation^25^. By binding and opening ME-specific regulatory regions, EOMES promotes chromatin accessibility and facilitates the activation of lineage-specific gene programs, in cooperation with the SWI/SNF chromatin-remodeling complex^25^. Based on this function as a chromatin remodeler, we hypothesize that EOMES acts in a similar manner in the cerebellum to govern the transcriptional dynamics associated with UBC migration and maturation through the remodeling of chromatin states. EOMES may enable the cooperative action of critical UBC TFs, such as LMX1A and SOX4, thereby promoting the activation of GRNs necessary for UBC development and maturation.

Overall, our study establishes that early human and mouse RL development is highly conserved, sharing a common molecular organization and TF co-expression programs that define UBC lineage states. Building on this conserved developmental framework, we show that *Eomes* is not a determinant of UBC cell fate as previously assumed, but instead functions as a developmental gatekeeper that enables differentiating UBCs to exit the RL, migrate appropriately, and complete their maturation by orchestrating downstream UBC transcriptional networks, including the activity of the key regulator *Lmx1a*.

## MATERIALS AND METHODS

### Key resources table

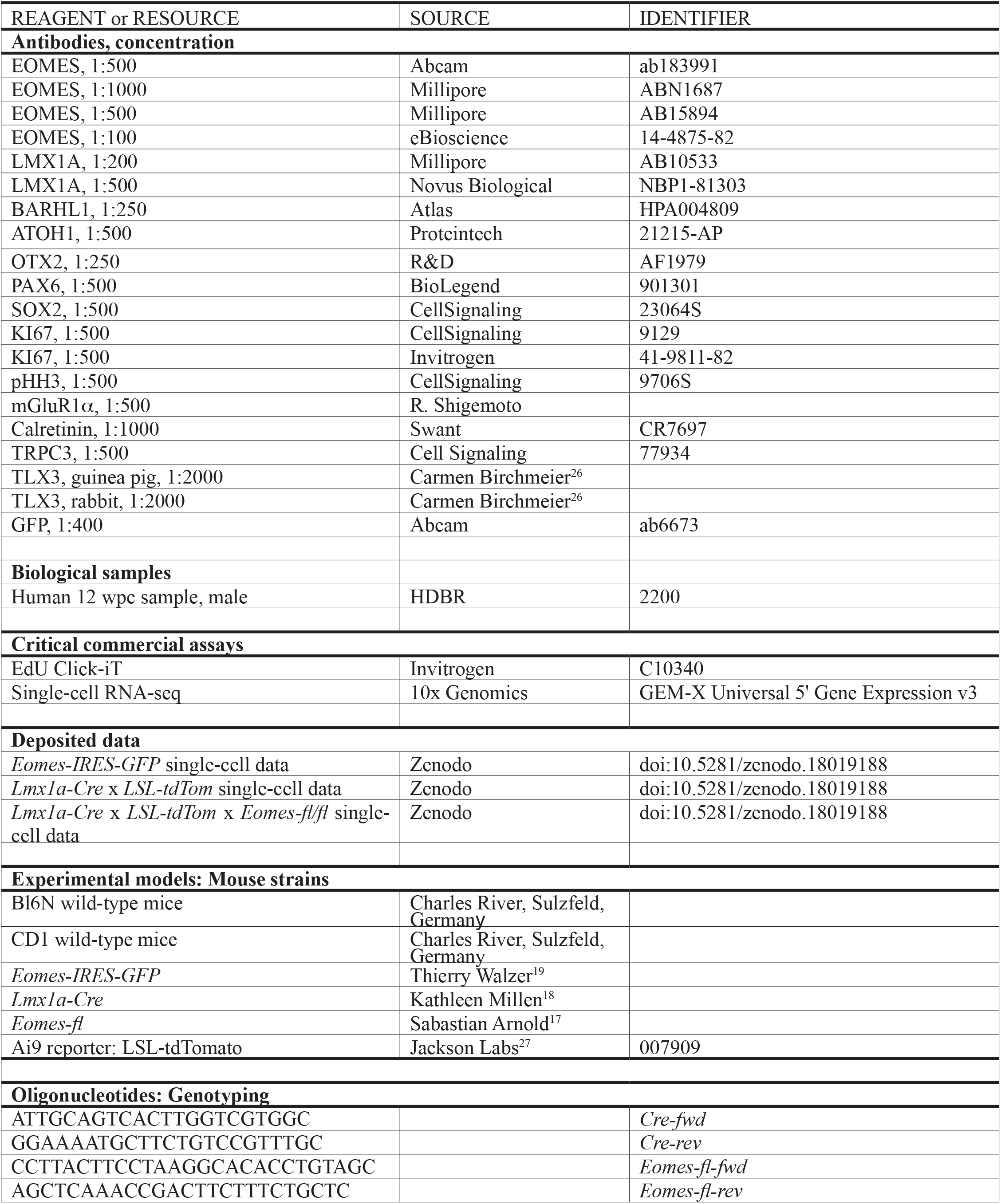
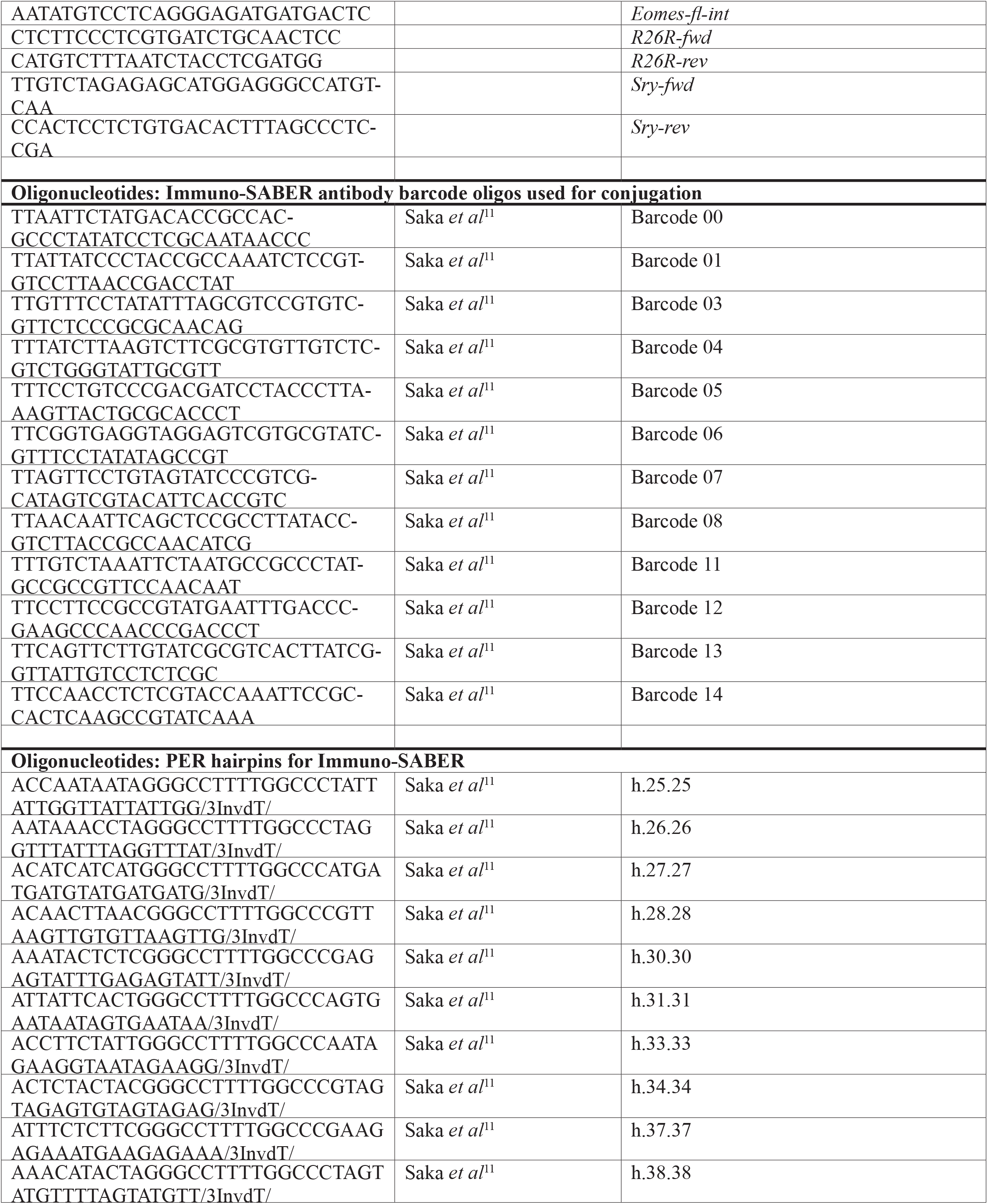
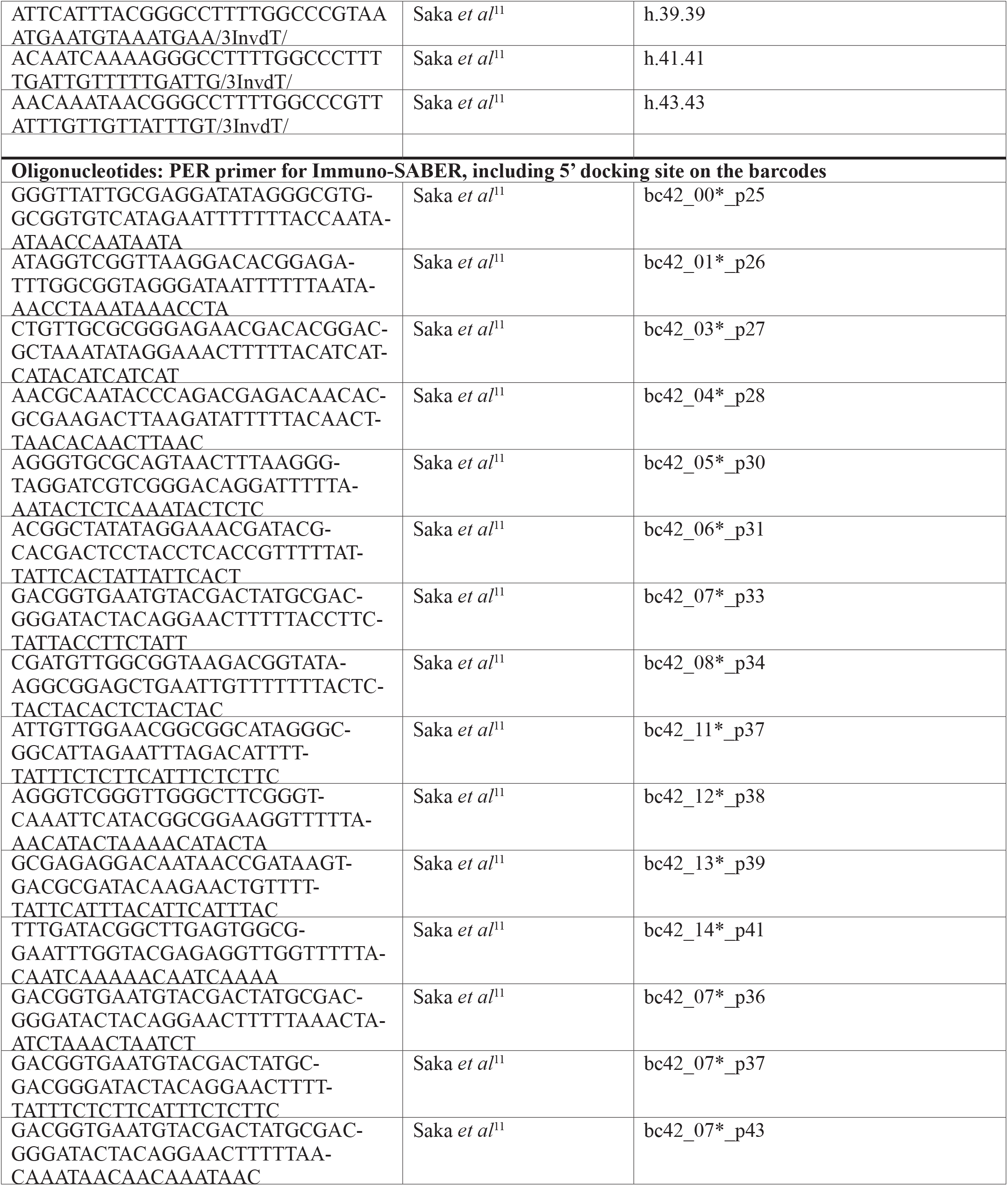
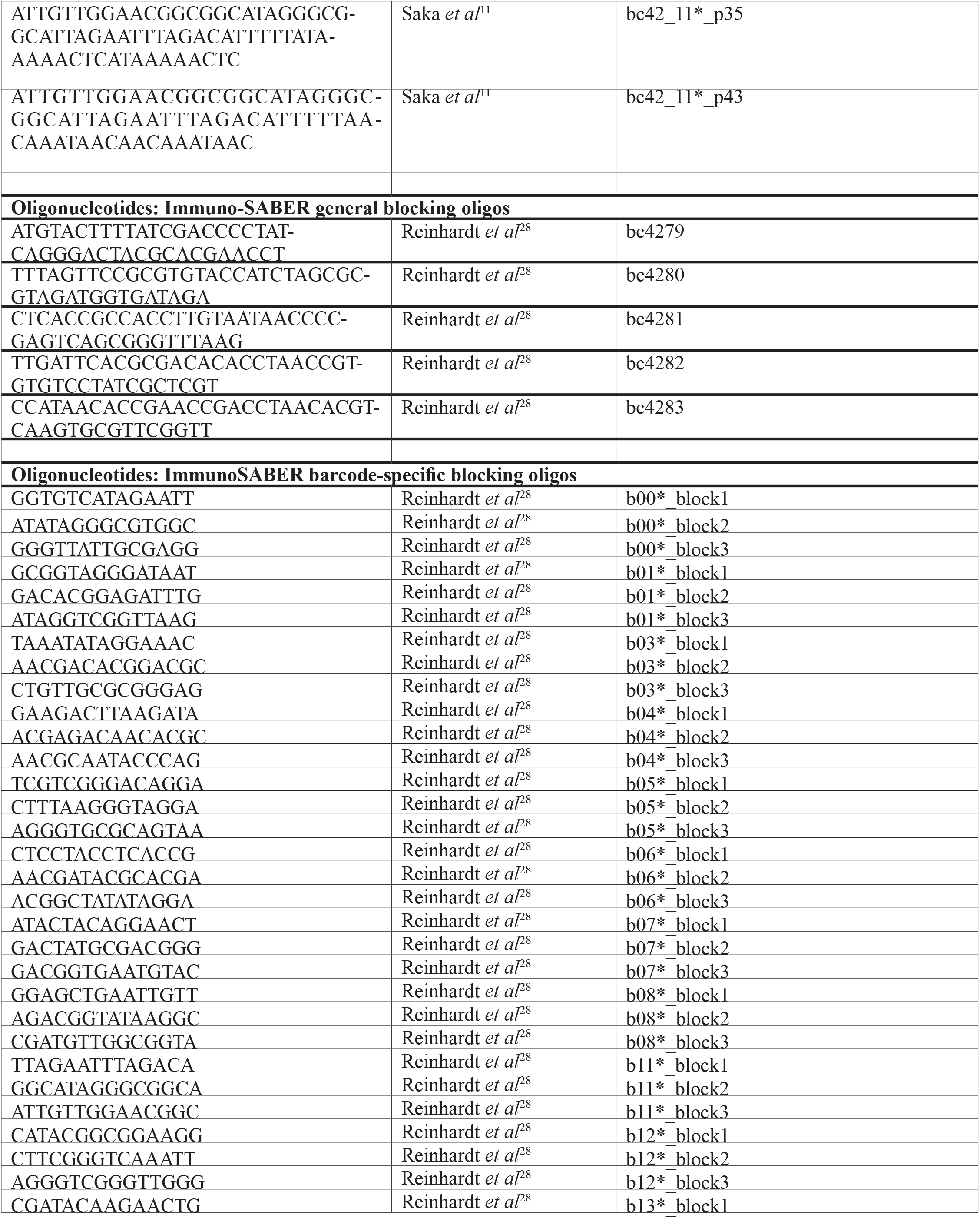
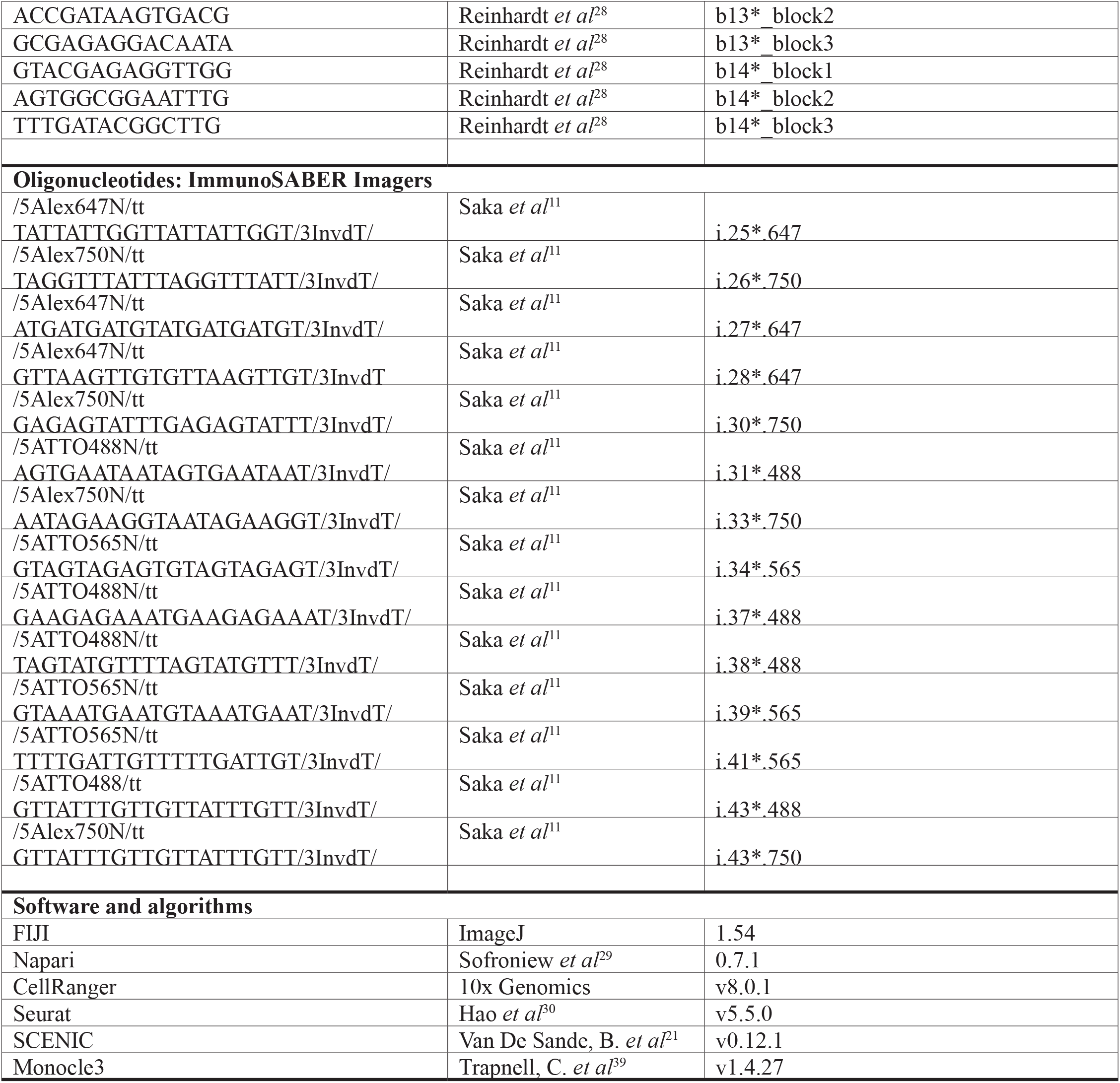

## EXPERIMENTAL MODEL DETAILS

### Animals

Animal strains are listed in the Key Resources Table. The animals were housed under a 12h/12h dark/light cycle controlled for temperature (20-24°C) and humidity (40-65%) with *ad libitum* access to food and water. Genetically engineered mice were bred on a C57BL/6N background. Animal experiments were approved by the responsible local authorities: Regierungspräsidium Karlsruhe (G-69/21) and the DKFZ Central Animal Laboratory (DKFZ383, DKFZ443-24). The *Lmx1a-Cre* transgene is integrated on the X-chromosome, so 50% of the cells in female animals undergo X-linked inactivation^31^. Therefore, we examined male *Lmx1a-Cre* animals. The morning of vaginal plug was designated as E0, and the day of birth as P0. We imaged at the midline vermis because of relationship of this area to the localization of Group 3/4 medulloblastoma and the site of Dandy-Walker Syndrome.

### EdU injections

E16 pregnant dams or P0 pups were injected intraperitoneally with 10 mg/ml stock solution of EdU (50 mg/kg, dissolved in PBS, Invitrogen). Mice were sacrificed 2h post-injection.

### Genotyping

Genotyping primer sequences are available in Key Resource Table. Tissue samples were either sent to Transnetyx for genotyping or performed in-house. For in-house genotyping, tissue was incubated overnight at 55°C in One-Step Tail Buffer (100 mM Tris-Cl, pH 8.3, 200 mM NaCl, 5 mM EDTA, 1% tergitol, 5 mg/ml ProteinaseK), followed by 1h heat-inactivation at 85°C. PCRs were performed with GoTaq.

### Human fetal sample

The human prenatal sample was donated voluntarily to the MRC Wellcome Trust Human Developmental Biology Resource (HDBR; UK) by women who had an elective abortion and had given written informed consent to donate fetal tissues for research. The use of human samples was approved by an ethics committee in Heidelberg (authorization S-220/2017, S-417/2025), North East-Newcastle & North Tyneside (REC reference 18/NE/0290), London-Fulham (REC reference 18/LO/0822).

## METHODS DETAILS

### Immuno-SABER

Immuno-SABER was performed as described previously^28^, without decoy imagers. Briefly, 11 rabbit antibodies were individually conjugated with a specific 42 nucleotide barcode using oYo-Link® Conjugation Technology (AlphaThera). 1 μg of primary antibody was mixed with 1 μl of oYo-Link® barcode and placed on ice for 2h under UV light in the cold room. Conjugated antibodies were stored in the same conditions as before conjugation and were aliquoted into minimum 5 μl aliquots to prevent multiple freeze-thaw cycles.

To amplify the signal of each primary antibody, extending single-stranded oligos were produced using primer exchange reactions (PERs), as described previously^11^. PER primers were ordered from Sigma using desalted purification. PER hairpins were ordered from IDT as 100 nmol, HPLC purified. A primer-hairpin combination specific for each barcode was mixed in PBS with 10 mM MgSO_4_, 500 U/ml Bst LF polymerase (McLab), 300 μM of dATP, dCTP and dTTP each (Carl Roth), 100 nM of Clean.G hairpin, and 0.1-0.6 μM hairpins, obtaining 600-750 nucleotide amplifier concatemers. PERs were then purified using the MinElute Gel Extraction Kit (QIAGEN). For staining, PFA/cryo slides were washed with 0.3% TBS-T (Triton X-100). Blocking and permeabilization was performed using PBS supplemented with 2% BSA, 0.3% Triton X-100, 0.1% Dextran sulfate, 4 mM EDTA, 0.5 mg/ml sheared salmon sperm and 800 nM of a blocking oligo pool^28^. Blocking Buffer was incubated for minimum of 2h. Oligo-conjugated primary antibodies were incubated overnight at 4°C in a humidified chamber. After washing, antibodies were fixed to their antigens with 5 mM Bis(sulfosuccinimidyl) suberate in PBS for 30 min followed by quenching with 100 mM NH_4_Cl for 10 min. Incubation of PERs was performed for 1h in a humidified chamber at 37°C.

Imagers were diluted to 1 μM in 0.1% PBS-T and incubated for 30 min in the dark at RT. Slides were mounted using Slow Fade Antifade Gold (Invitrogen). Cover slips were secured with FixoGum. Imagers without primary antibodies did not produce any signal. The procedure was completed with subsequent washing and imaging rounds. On the fourth day, traditional secondary antibody staining against goat and rat, was performed to label OTX2 and SOX2, respectively. All images were acquired on the Leica Thunder with the Leica-K8-A22M726030 camera and 40× Oil Immersion Objective (HC PL APO CS2 40×/1.30 Oil UV). Images were acquired from 3 animals per time points, analyzing 1 section per animal.

### Immuno-SABER imaging processing and analysis

The acquired Leica images were stored in the .*lif* format and processed using a custom Python pipeline. Specifically, *aicsimageio* was used to lazily load the volumetric arrays^32^. To identify the most-in-focus optical plane of each section, the variance of the Laplacian operator was calculated for each optical plane of the DAPI image of each imaging round using the *skimage*.*filters*.*laplace* implementation^34^ with a default kernel size of three for both the *x* and *y* dimension. For each DAPI image, the optical plane having the highest variance of the Laplacian operator was deemed to be the sharpest optical plane and extracted from the volumetric array. *Matplotlib*^36^ was used to create line plots with the x axis as the optical planes for a specific DAPI image and the y-axis as the measure of sharpness These were inspected to assess the focus behavior and served as an initial validation of the automated plane selection. Subsequently, for each other channel image in the same imaging round, the same optical plane was extracted and combined with the extracted DAPI image to create a 2D multichannel image.

For spatial alignment of the imaging rounds of each tissue section, rigid-body or affine registration was performed using *pystackreg*^38^, a Python implementation of the *FIJI* plugin *TurboReg*. The required transformations for Materials and Methods for spatial alignment were estimated using the DAPI channel of one imaging round as reference image and learning the transformation of the DAPI channels of the other imaging rounds to that reference image. Subsequently, the learned transformation was applied to the DAPI channel and each other channel image of the same imaging round. The resulting 16-bit 2D images were stored using the *ome-tiff* format^40^. Next, the multichannel images of each section were jointly visualized in *napari*^29^ to assess matching cell content of the extracted optical planes and spatial alignment. Sections showing improper alignment or non-matching cell content between imaging rounds were not subsequently processed and analyzed.

The image channels of each section corresponding to markers of interest were then segmented using *Cellpose*^42^. More specifically the *cyto3* model was used as a baseline model and iteratively trained by feeding in *napari* corrected segmentations to the model. This improved the segmentation quality over time. Only those cells positive for the specific marker were segmented, resulting in a segmentation map of marker positive cells for each marker of interest. Each segmentation map was overlaid in *napari* on the corresponding channel image and assessed for segmentation quality by two researchers. Lastly, the segmentation maps were used to obtain a count of cells per anatomical region with the marker profiles of interest.

Statistical analysis was performed using Graph Pad Prism (v10.5.0) using Ordinary one-way ANOVA or T-test as appropriate, with a p<0.05 considered as statistically significant.

### Histology and immunostaining

For histology, samples of murine brains were formalin-fixed, paraffin-embedded, and sectioned at 5 μm. The sections were stained with hematoxylin and eosin (H&E), and imaged on a Histoscanner PrimeHisto XE (Pacific Image Electronics Co). Immunostaining was performed as described previously^33^. Brains were harvested, fixed overnight in 4% paraformaldehyde (PFA) at 4°C, soaked in 30% sucrose in PBS overnight, and embedded in optimal cutting temperature (OCT) compound. Frozen 10-μm-thick sections were collected on Fisher Superfrost Plus slides using a cryostat. Sections were blocked for 30 min at roomRT with 10% normal donkey serum and incubated with primary antibody overnight at 4°C (see Key Resources Table). After washing with PBST, sections were incubated with appropriate fluorescence secondary antibodies for 1h at RT. Images were acquired using a widefield Zeiss Cell Observer or confocal laser-scanning microscopes (Observer.Z1, LSM710 ConfoCor3 and LSM780, Zeiss). For quantifications using FIJI, 3 animals were analyzed per genotype per time point, counting 3-4 sections per animal, blinded to genotype.

### Single-cell RNA-sequencing: Sample preparation and library construction

*Eomes-IRES-GFP* homozygous knock-in male animals were paired with CD1 outbred females to increase brood size. *Lmx1a-Cre*; *Eomes-fl/fl*; Ai9 and *Lmx1a-Cre*; Ai9 animals were bred on a Bl6N background. At embryonic day E16 or at the day of birth (P0), the cerebellum was dissected from the rest of the brain. The meninges were removed, and the cerebella were collected in Hibernate A media supplemented with B27+RA (2%), Glutamax (200 mM) and Gentamicin. Processing of the tissue was done using the Papain Dissociation System from Worthington (LK003153). Extracted single cells were prepared for FACS by concentrating to 2-10x10^6^ cells/ml in FACS buffer (1x PBS ,1% BSA). FACS was performed using a BD FACSAria III Cell Sorter. In the *Eomes-IRES-GFP* samples, GFP+ singlets were collected, whereas in the *Lmx1a-Cre; Eomes-fl; Ai9 and Lmx1a-Cre; Ai9* animals, tdTom+ singlets were isolated. A total of 20,000 sorted cells were then processed with the 10X Genomics Chromium Next GEM Single Cell 3’ Reagent Kits v3.1 (Dual Index), according to the manufacturer’s instructions. Library preparation was followed by sequencing on an Illumina NextSeq™ 1000/2000.

### Initial data pre-processing, quality control (QC), and object generation

De-multiplexed reads were aligned with *CellRanger* (v8.0.1) to the mouse genome assembly GRCm39 (2024-A, reference dataset from 10X Genomics). Raw unique molecular identifier (UMI) count matrices were imported from the 10x Genomics output file (raw_feature_bc_matrix. h5) using *Read10X_h5* and converted into a Seurat object, initially retaining only barcodes with ≥3 detected cells and ≥200 expressed genes.

To assess data quality, mitochondrial-encoded genes were identified for each cell, and the proportion of mitochondrial UMIs was calculated relative to the total transcript counts. A metadata table was generated summarizing, for each cell, the number of detected genes (nFeature_RNA), total UMI counts (nCount_RNA), and mitochondrial fraction. A minimum cutoff of 750 UMIs and at least 500 detected genes per cell was applied, and cells with >5% mitochondrial UMIs Materials and Methods for were excluded. All cells meeting these three criteria (≥500 genes, ≥750 UMIs, ≤5% mitochondrial reads) were labeled, extracted by barcode, and used to generate a filtered Seurat object with *Seurat v5*.*3*.*1. DoubletFinder* was then used to identify and remove doublets prior to downstream analyses.

### Normalization, scaling and dimensionality reduction

The filtered dataset was normalized using the global-scaling method *NormalizeData (LogNormalize)*, followed by *ScaleData*. Feature selection was performed with *FindVariableFeatures*, retaining 5,000 features per dataset. Dimensionality reduction was conducted using *RunPCA*, and UMAP embeddings were computed from principal components selected according to their distribution in the *ElbowPlot*.

### Clustering, visualization, and annotation

For clustering, resolution and k-nearest-neighbor parameters were tested with *FindNeighbors*. Cluster identities were assigned by examining marker gene expression and guided by the E16 and P0 Immuno-SABER stainings. Populations containing clusters with large numbers of cells were subsetted and reclustered to resolve biologically meaningful subclusters. Additional quality controls for nFeature_RNA, total UMI counts (nCount_RNA), and mitochondrial fraction were performed after clustering and the clusters with poor QC content were filtered out.

### Differentially expressed gene (DEG) analysis

For differential expression across annotated lineages and between control and mutant-specific clusters, cells of interest were first pseudobulked based on their genotype. Differential expression was performed between relevant subsets using Seurat’s *FindMarkers* with the Wilcoxon test. No pre-filtering thresholds were used for minimum expressing fraction or log-fold-change (min.pct = 0, logfc. threshold = 0), allowing all genes to be tested. Significant genes were then defined by adjusted p<0.05 and |average log_2_FC|≥1. DEG results were visualized with a volcano plot to highlight genes exhibiting both large fold-changes and strong statistical support.

### Gene Set Enrichment Analysis (GSEA) and Gene Ontology (GO) Terms

For each genotype and timepoint, the DEGs were first ranked based on the signed average log_2_FC x -log_10_(adjusted p-value). The pre-ranked DEGs were prepared in a clean format where gene symbols were converted to gene names, the duplicate genes and NA values were removed from the list, and the cleaned DEGs were sorted in a descending order. GSEA^35^ was run for *Biological Process (BP)* and top pathways with the highest Normalized Enrichment Score (NES) and significant p-values (p<0.05) were visualized between UBC and UBC cKO for each time point. GO terms^37^ were also analyzed on the selected DEGs with the adjusted p<0.05 and |average log_2_FC|≥1 for *BP* separately for genotype and timepoint.

### Pseudotime trajectory analysis

To infer the trajectory, the cells belonging to the UBC lineage in the joint filtered Seurat object were first subset and the workflow for *monocle3*^39^ was followed for this analysis. The subset Seurat object was then converted into a CDS object (*as*.*cell_data_set*). Next, cells were grouped into clusters and partitions (*cluster_cells*), which are disconnected in the reduced dimensional space and the trajectory was inferred (*learn_graph*). The root cells were defined as the clusters belonging to “GC_UBC progenitors” and then each individual cell was ordered along the predicted trajectory (*order_cells*).

### Gene regulatory network inference

Single Cell Regulatory Network Inference and Clustering (SCENIC^21^) workflow was applied via *pyscenic*^41^ on the single cell RNA-seq dataset for *Lmx1a-Cre*; *Eomes-fl/fl*; Ai9 and *Lmx1a-Cre*; Ai9 animals separately to infer the gene regulatory network differences upon *Eomes* cKO. First, the potential targets for each TF in the dataset were identified from the gene count matrix by *GRNBoost2*^43^ *(grn)* based on the calculated correlations between the gene expressions and individual cells. Then, the indirect targets were eliminated by using the *cis*-regulatory motif analysis *(ctx)*, which allowed to obtain the refined TF-direct targets (regulons). Finally as the last step of *pyscenic* workflow, regulon activity in individual cells was calculated by *aucell* scoring.

## Author Contributions

V.A., L.M.S, P.B.G.d.S, A.S., A.S.D. and L.M.K. performed experiments and analyzed data. W-M.A.M.V. designed and implemented high-content imaging analysis code. V.A., P.J. and M.S. analyzed single-cell RNA-seq data. L.S., J.N., R.R., F.S., N.H. performed experiments. A.S.D., S.M.P., M.Z., and H.K. advised on the experimental design and provided critical data and reagents. S.K.S., and L.M.K. supervised the study. L.M.K. designed the study. V.A., L.M.S., and L.M.K. drafted the first version of the manuscript. All authors provided feedback and approved its final version.

## Data and code availability

Raw single-cell sequencing data deposited to Zenodo under doi:10.5281/zenodo.18019188. Scripts and processed single-cell objects, including normalized counts, metadata, and clustering information are available at github.com/kutscher-lab/UBC-Development-Analysis-Pipeline and Zenodo.

## Acknowledgments

We acknowledge the access and services provided by the Imaging Centre at the European Molecular Biology Laboratory (EMBL IC), generously supported by the Boehringer Ingelheim Foundation. We thank Dieter Walsh for experimental advice and guidance concerning the high-content imaging experiments. We acknowledge the Joint MRC/Wellcome (MR/R006237/1) Human Developmental Biology Resource, especially Steven Lisgo. We thank Ryuichi Shigemoto for sharing the mGluR1α antibody, Robert Hevner and Ray Daza for sharing antibodies and knowledge, and Carmen Birchmeier for TLX3 antibodies. We thank Kathleen Millen for the *Lmx1a-Cre* mouse line, Sebastian Arnold for the *Eomes-fl* mouse line, and Thierry Walzer for the *EOMES-IRES-GFP* mouse line. We gratefully acknowledge the following core facilities at the DKFZ: Genomics core facility, Single-cell open lab, light microscopy core facility, and the Center for Preclinical Studies. We thank members of the Kutscher Lab for critical reading of the manuscript and members of the Saka Lab for technical support with the Immuno-SABER assay. We thank Annarita Patrizi for critical review of the manuscript. Leonie Ringrose (Science Kitchen) provided scientific editing. Funding was provided by the Emmy Noether Programme from the German Research Foundation to L.M.K. (551030459), the European Research Council to S.M.P. (819894), and Fight Kids Cancer to L.M.K., S.M.P., and A.S.D. (Fight4MB). S.K.S. was supported by core funding from European Molecular Biology Laboratory. W-M.A.M.V. acknowledges financial support from Cellzome, a GSK company. M.S. was supported by the Simons Foundation Autism Research Initiative (SFARI) Bridge to Independence Award (SFI-AN-AR-Independence Postdoctoral-00007139. Some figure panels were created in BioRender. Figure 2E: Akçay, V. (2026) https://BioRender.com/q5jnrxx. Figure 3A: Akçay, V. (2026) https://BioRender.com/0p4xt2u

## References

1. Diedrichsen, J., and McDougle, S.D. (2026). How does the cerebellum contribute to cognitive functions? PLoS Biol 24, e3003688. 10.1371/journal.pbio.3003688.

2. Haldipur, P., Millen, K.J., and Aldinger, K.A. (2022). Human Cerebellar Development and Transcriptomics: Implications for Neurodevelopmental Disorders. Annu Rev Neurosci 45, 515–531. 10.1146/annurev-neuro-111020-091953.

3. Okonechnikov, K., Joshi, P., Sepp, M., Leiss, K., Sarropoulos, I., Murat, F., Sill, M., Beck, P., Chan, K.C.-H., Korshunov, A., et al. (2023). Mapping pediatric brain tumors to their origins in the developing cerebellum. Neuro-Oncology, noad124. 10.1093/neuonc/noad124.

4. Hendrikse, L.D., Haldipur, P., Saulnier, O., Millman, J., Sjoboen, A.H., Erickson, A.W., Ong, W., Gordon, V., Coudière-Morrison, L., Mercier, A.L., et al. (2022). Failure of human rhombic lip differentiation underlies medulloblastoma formation. Nature 609, 1021–1028. 10.1038/s41586-022-05215-w.

5. Smith, K.S., Bihannic, L., Gudenas, B.L., Haldipur, P., Tao, R., Gao, Q., Li, Y., Aldinger, K.A., Iskusnykh, I.Y., Chizhikov, V.V., et al. (2022). Unified rhombic lip origins of group 3 and group 4 medulloblastoma. Nature 609, 1012–1020. 10.1038/s41586-022-05208-9.

6. Haldipur, P., Aldinger, K.A., Bernardo, S., Deng, M., Timms, A.E., Overman, L.M., Winter, C., Lisgo, S.N., Razavi Silvestri, E., et al. (2019). Spatiotemporal expansion of primary progenitor zones in the developing human cerebellum. Science 366, 454–460. 10.1126/science.aax7526.

7. Yeung, J., Ha, T.J., Swanson, D.J., Choi, K., Tong, Y., and Goldowitz, D. (2014). Wls Provides a New Compartmental View of the Rhombic Lip in Mouse Cerebellar Development. Journal of Neuroscience 34, 12527–12537. 10.1523/JNEUROSCI.1330-14.2014.

8. Phoenix, T.N. (2022). The origins of medulloblastoma tumours in humans. Nature 609, 901–903. 10.1038/d41586-022-02951-x.

9. Butts, J.C., Wu, S.-R., Durham, M.A., Dhindsa, R.S., Revelli, J.-P., Ljungberg, M.C., Saulnier, O., McLaren, M.E., Taylor, M.D., and Zoghbi, H.Y. (2024). A single-cell transcriptomic map of the developing Atoh1 lineage identifies neural fate decisions and neuronal diversity in the hindbrain. Developmental Cell 59, 2171–2188.e7. 10.1016/j.dev-cel.2024.07.007.

10. McDonough, A., Elsen, G.E., Daza, R.M., Bachleda, A.R., Pizzo, D., DelleTorri, O.M., and Hevner, R.F. (2021). Unipolar (Dendritic) Brush Cells Are Morphologically Complex and Require Tbr2 for Differentiation and Migration. Front. Neurosci. 14, 598548. 10.3389/fnins.2020.598548.

11. Saka, S.K., Wang, Y., Kishi, J.Y., Zhu, A., Zeng, Y., Xie, W., Kirli, K., Yapp, C., Cicconet, M., Beliveau, B.J., et al. (2019). Immuno-SABER enables highly multiplexed and amplified protein imaging in tissues. Nat Biotechnol 37, 1080–1090. 10.1038/s41587-019-0207-y.

12. Sepp, M., Leiss, K., Murat, F., Okonechnikov, K., Joshi, P., Leushkin, E., Spänig, L., Mbengue, N., Schneider, C., Schmidt, J., et al. (2024). Cellular development and evolution of the mammalian cerebellum. Nature 625, 788–796. 10.1038/s41586-023-06884-x.

13. Visvanathan, A., Saulnier, O., Chen, C., Haldipur, P., Orisme, W., Delaidelli, A., Shin, S., Millman, J., Bryant, A., Abeysundara, N., et al. (2024). Early rhombic lip Protogenin+ve stem cells in a human-specific neurovascular niche initiate and maintain group 3 medulloblastoma. Cell 187, 4733–4750.e26. 10.1016/j.cell.2024.06.011.

14. Li, Z., Tyler, W.A., Zeldich, E., Santpere Baró, G., Okamoto, M., Gao, T., Li, M., Sestan, N., and Haydar, T.F. (2020). Transcriptional priming as a conserved mechanism of lineage diversification in the developing mouse and human neocortex. Sci Adv 6, eabd2068. 10.1126/sciadv.abd2068.

15. Krishnamurthy, A., Lee, A.S., Bayin, N.S., Stephen, D.N., Nasef, O., Lao, Z., and Joyner, A.L. (2024). Engrailed transcription factors direct excitatory cerebellar neuron diversity and survival. Development 151, dev202502. 10.1242/dev.202502.

16. Englund, C. (2006). Unipolar Brush Cells of the Cerebellum Are Produced in the Rhombic Lip and Migrate through Developing White Matter. Journal of Neuroscience 26, 9184–9195. 10.1523/JNEUROSCI.1610-06.2006.

17. Arnold, S.J., Hofmann, U.K., Bikoff, E.K., and Robertson, E.J. (2008). Pivotal roles for eomesodermin during axis formation,epithelium-to-mesenchyme transition and endoderm specification in the mouse. Development 135, 501–511. 10.1242/dev.014357.

18. Chizhikov, V.V., Lindgren, A.G., Mishima, Y., Roberts, R.W., Aldinger, K.A., Miesegaes, G.R., Currle, D.S., Monuki, E.S., and Millen, K.J. (2010). Lmx1a regulates fates and location of cells originating from the cerebellar rhombic lip and telencephalic cortical hem. Proceedings of the National Academy of Sciences 107, 10725–10730. 10.1073/pnas.0910786107.

19. Daussy, C., Faure, F., Mayol, K., Viel, S., Gasteiger, G., Charrier, E., Bienvenu, J., Henry, T., Debien, E., Hasan, U.A., et al. (2014). T-bet and Eomes instruct the development of two distinct natural killer cell lineages in the liver and in the bone marrow. J Exp Med 211, 563–577. 10.1084/jem.20131560.

20. Miyazawa, K., Himi, T., Garcia, V., Yamagishi, H., Sato, S., and Ishizaki, Y. (2000). A Role for p27/Kip1 in the Control of Cerebellar Granule Cell Precursor Proliferation. J. Neurosci. 20, 5756–5763. 10.1523/JNEUROS-CI.20-15-05756.2000.

21. Van De Sande, B., Flerin, C., Davie, K., De Waegeneer, M., Hulselmans, G., Aibar, S., Seurinck, R., Saelens, W., Cannoodt, R., Rouchon, Q., et al. (2020). A scalable SCENIC workflow for single-cell gene regulatory network analysis. Nat Protoc 15, 2247–2276. 10.1038/s41596-020-0336-2.

22. Northcott, P.A., Buchhalter, I., Morrissy, A.S., Hovestadt, V., Weischenfeldt, J., Ehrenberger, T., Gröbner, S., Segura-Wang, M., Zichner, T., Rudneva, V.A., et al. (2017). The whole-genome landscape of medulloblastoma subtypes. Nature 547, 311–317. 10.1038/nature22973.

23. Jones, D.T.W., Jäger, N., Kool, M., Zichner, T., Hutter, B., Sultan, M., Cho, Y.-J., Pugh, T.J., Hovestadt, V., Stütz, A.M., et al. (2012). Dissecting the genomic complexity underlying medulloblastoma. Nature 488, 100–105. 10.1038/nature11284.

24. Gilthorpe, J.D., Papantoniou, E.-K., Chédotal, A., Lumsden, A., and Wingate, R.J.T. (2002). The migration of cerebellar rhombic lip derivatives. Development 129, 4719–4728. 10.1242/dev.129.20.4719.

25. Schröder, C.M., Zissel, L., Mersiowsky, S.-L., Tekman, M., Probst, S., Schüle, K.M., Preissl, S., Schilling, O., Timmers, H.Th.M., and Arnold, S.J. (2025). EOMES establishes mesoderm and endoderm differentiation potential through SWI/SNF-mediated global enhancer remodeling. Developmental Cell 60, 735–748.e5. 10.1016/j.devcel.2024.11.014.

26. Müller, T., Anlag, K., Wildner, H., Britsch, S., Treier, M., and Birchmeier, C. (2005). The bHLH factor Olig3 coordinates the specification of dorsal neurons in the spinal cord. Genes Dev. 19, 733–743. 10.1101/gad.326105.

27. Madisen, L., Zwingman, T.A., Sunkin, S.M., Oh, S.W., Zariwala, H.A., Gu, H., Ng, L.L., Palmiter, R.D., Hawrylycz, M.J., Jones, A.R., et al. (2010). A robust and high-throughput Cre reporting and characterization system for the whole mouse brain. Nat Neurosci 13, 133–140. 10.1038/nn.2467.

28. Reinhardt, R., Straka, T., Vierdag, W.-M., Jevdokimenko, K., Hecht, F., Pianfetti, E., Hudelmaier, T., Lai, H., Fouquet, W., Fahrbach, F., et al. (2026). Enabling high-plex spectral imaging via DNA-barcoded signal tuning and panel optimization. Preprint, 10.64898/2026.03.18.709053 https://doi.org/10.64898/2026.03.18.709053.

29. Sofroniew, N., Lambert, T., Bokota, G., Nunez-Iglesias, J., Sobolewski, P., Sweet, A., Gaifas, L., Evans, K., Burt, A., Doncila Pop, D., et al. (2026). napari: a multi-dimensional image viewer for Python. Version v0.7.0 (Zenodo). 10.5281/ZENODO.19183461 https://doi.org/10.5281/ZE-NODO.19183461.

30. Hao, Y., Stuart, T., Kowalski, M.H., Choudhary, S., Hoffman, P., Hartman, A., Srivastava, A., Molla, G., Madad, S., Fernandez-Granda, C., et al. (2024). Dictionary learning for integrative, multimodal and scalable single-cell analysis. Nat Biotechnol 42, 293–304. 10.1038/s41587-023-01767-y.

31. Parichha, A., Suresh, V., Chatterjee, M., Kshirsagar, A., Ben-Reuven, L., Olender, T., Taketo, M.M., Radosevic, V., Bobic-Rasonja, M., Trnski, S., et al. (2022). Constitutive activation of canonical Wnt signaling disrupts choroid plexus epithelial fate. Nat Commun 13, 633. 10.1038/s41467-021-27602-z.

32. Sherman, J., Toloudis Swain-Bowden, M., Talley Lambert, LeRoy, S., Brown, E.M., BrianWhitneyAI, Matte Bailey, Basu Chaudhuri, Johnson, G., et al. (2023). AllenCellModeling/aicsimageio: OME metadata for ND2 reader and ome-types upgrade. Version v4.12.1 (Zenodo). 10.5281/ZE-NODO.4906608 https://doi.org/10.5281/ZENODO.4906608.

33. Arnskötter, F., da Silva, P.B.G., Schouw, M.E., Lukasch, C., Bianchini, L., Sieber, L., Garcia-Lopez, J., Ahmad, S.T., Li, Y., Lin, H., et al. (2024). Loss of Elp1 in cerebellar granule cell progenitors models ataxia phenotype of Familial Dysautonomia. Neurobiol Dis 199, 106600. 10.1016/j.nbd.2024.106600.

34. Van Der Walt, S., Schönberger, J.L., Nunez-Iglesias, J., Boulogne, F., Warner, J.D., Yager, N., Gouillart, E., and Yu, T. (2014). scikit-image: image processing in Python. PeerJ 2, e453. 10.7717/peerj.453.

35. Subramanian, A., Tamayo, P., Mootha, V.K., Mukherjee, S., Ebert, B.L., Gillette, M.A., Paulovich, A., Pomeroy, S.L., Golub, T.R., Lander, E.S., et al. (2005). Gene set enrichment analysis: A knowledge-based approach for interpreting genome-wide expression profiles. Proc. Natl. Acad. Sci. U.S.A. 102, 15545–15550. 10.1073/pnas.0506580102.

36. Hunter, J.D. (2007). Matplotlib: A 2D Graphics Environment. Comput. Sci. Eng. 9, 90–95. 10.1109/MCSE.2007.55.

37. Gene Ontology Consortium (2004). The Gene Ontology (GO) database and informatics resource. Nucleic Acids Research 32, 258D–261. 10.1093/nar/gkh036.

38. Thevenaz, P., Ruttimann, U.E., and Unser, M. (1998). A pyramid approach to subpixel registration based on intensity. IEEE Trans. on Image Process. 7, 27–41. 10.1109/83.650848.

39. Trapnell, C., Cacchiarelli, D., Grimsby, J., Pokharel, P., Li, S., Morse, M., Lennon, N.J., Livak, K.J., Mikkelsen, T.S., and Rinn, J.L. (2014). The dynamics and regulators of cell fate decisions are revealed by pseudotemporal ordering of single cells. Nat Biotechnol 32, 381–386. 10.1038/nbt.2859.

40. Besson, S., Leigh, R., Linkert, M., Allan, C., Burel, J.-M., Carroll, M., Gault, D., Gozim, R., Li, S., Lindner, D., et al. (2019). Bringing Open Data to Whole Slide Imaging. In Digital Pathology Lecture Notes in Computer Science., C. C. Reyes-Aldasoro, A. Janowczyk, M. Veta, P. Bankhead, and K. Sirinukunwattana, eds. (Springer International Publishing), pp. 3–10. 10.1007/978-3-030-23937-4_1.

41. Kumar, N., Mishra, B., Athar, M., and Mukhtar, S. (2021). Inference of Gene Regulatory Network from Single-Cell Transcriptomic Data Using pySCENIC. Methods Mol Biol 2328, 171–182. 10.1007/978-1-0716-1534-8_10.

42. Stringer, C., and Pachitariu, M. (2025). Cellpose3: one-click image restoration for improved cellular segmentation. Nat Methods 22, 592–599. 10.1038/s41592-025-02595-5.

43. Moerman, T., Aibar Santos, S., Bravo González-Blas, C., Simm, J., Moreau, Y., Aerts, J., and Aerts, S. (2019). GRN-Boost2 and Arboreto: efficient and scalable inference of gene regulatory networks. Bioinformatics 35, 2159–2161. 10.1093/bioinformatics/bty916.

