## Supplemental Figures for "Conserved cerebellar rhombic lip compartmentalization and *Eomes* regulatory networks govern unipolar brush cell development"

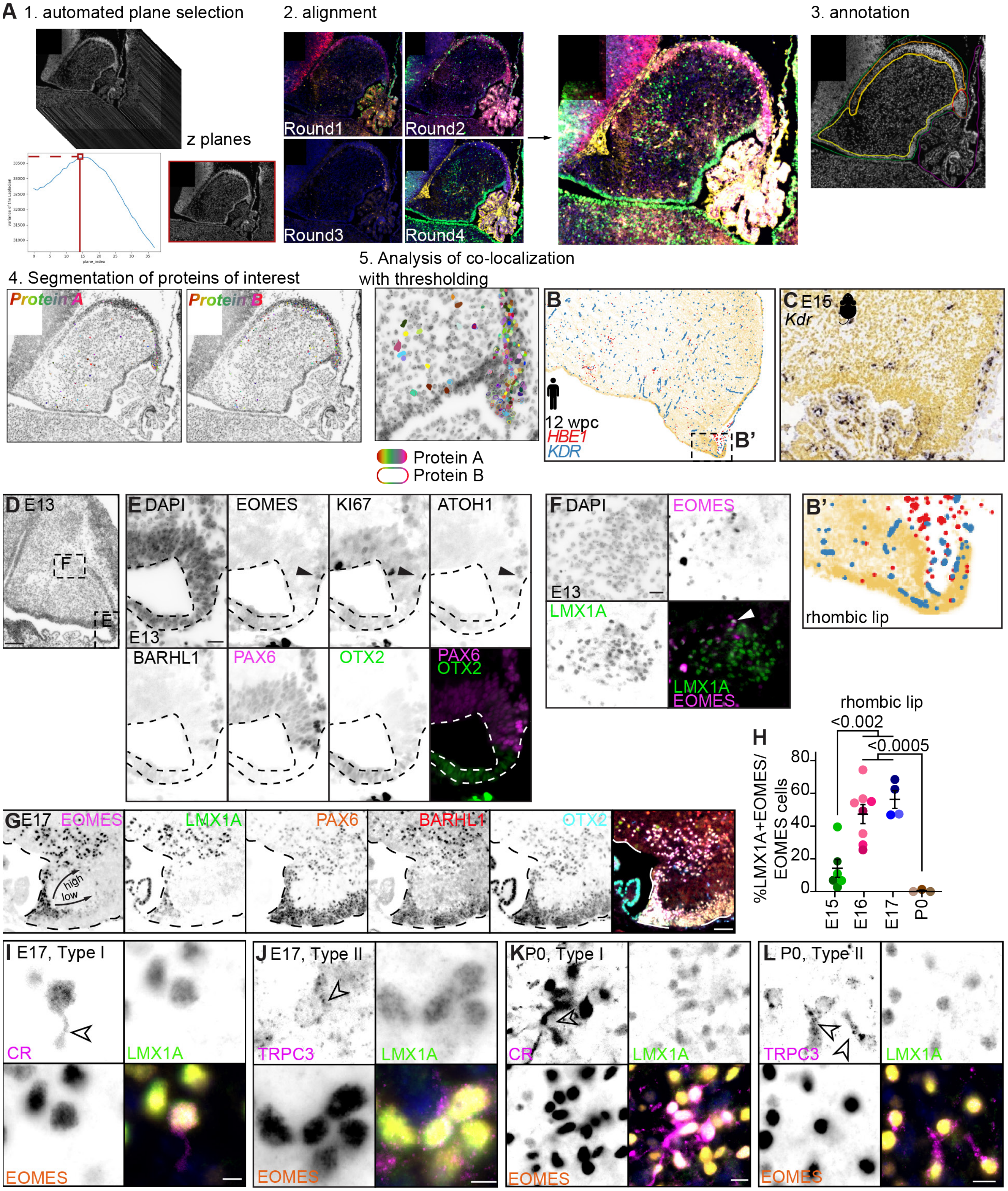


**Supplemental Figure S1** *legend next page* ►

◀*cont. from previous page*

**Figure S1**. **Immuno-SABER workflow and additional marker stainings**.

**A**) Immuno-SABER imaging analysis workflow: 1.) z-plane of interest from each round of imaging is selected automatically using peak variance of the Laplacian; 2.) Four rounds of imaging are aligned based on DAPI input to generate an aligned, composite image, where only the cerebellar anlage or specific lobes of the cerebellum are imaged; 3.) regions of interest are manually annotated; the focus of this paper is the cerebellar rhombic lip (red); 4.) Nuclei are segmented based on the protein of interest; 5.) co-localization of proteins of interest were analyzed after applying appropriate thresholds. **B**) Published spatial transcriptomic data[1] of the erythroid marker gene *HBE1* and the mural/endothelial marker gene *KDR* from adjacent 12 wpc section as shown in Figure 1G. B', zoom of inset shown in B. **C**) Published RNA *in situ* hybridization[2] of the mural/endothelial marker gene *Kdr* at E15 in murine midline cerebellum. **D**) DAPI staining of E13 wild-type cerebellar anlage. Scale bar, 100 µm. **E**) Zoom of the box in D. Immuno-SABER of indicated antibodies. Merged image shows OTX2:PAX6 boundary separating early choroid plexus from early rhombic lip. Black arrow, co-localization of triple positive EOMES, KI67, and ATOH1. Scale bar, 20 µm. **F**) Zoom of the box in D, Immuno-SABER of indicated antibodies. White arrow, EOMES and LMX1A co-localize in single NTZ cell. Scale bar, 20 µm. **G**) E17 rhombic lip Immuno-SABER of EOMES (magenta), LMX1A (green), PAX6 (orange), BARHL1 (red), and OTX2 (cyan). Arrows represent putative migration streams of high-expressing and lowly-expressing EOMES+ cells. Scale bar, 50 µm. **H**) Quantification of %LMX1A+EOMES positive cells within EOMES+ population in RL over time. Ordinary one-way ANOVA. Multiple comparisons revealed E15 and P0 as significantly different from E16 and E17, with displayed p-values. **I**) Immuno-SABER image of CR (magenta), LMX1A (green), and EOMES (orange), representing type I UBCs, at E17. Black arrowheads denote growing neurites. Scale bar, 5 µm. **J**) Immuno-SABER image of TRPC3 (magenta), LMX1A (green), and EOMES (orange), representing type II UBCs, at E17. Black arrowheads denote growing neurites. Scale bar, 5 µm. **K**) Immuno-SABER image of CR (magenta), LMX1A (green), and EOMES (orange), representing type I UBCs, at P0. Black arrowheads denote growing neurites. Scale bar, 10 µm. **L**) Immuno-SABER image of TRPC3 (magenta), LMX1A (green), and EOMES (orange), representing type II UBCs, at P0. Black arrowheads denote growing neurites. Scale bar, 10 µm.

### Supplemental References

1. Sepp, M. *et al.* Cellular development and evolution of the mammalian cerebellum. *Nature* **625**, 788–796 (2024).
2. Allen Developing Mouse Brain Atlas. Allen Institute for Brain Science. (2008).

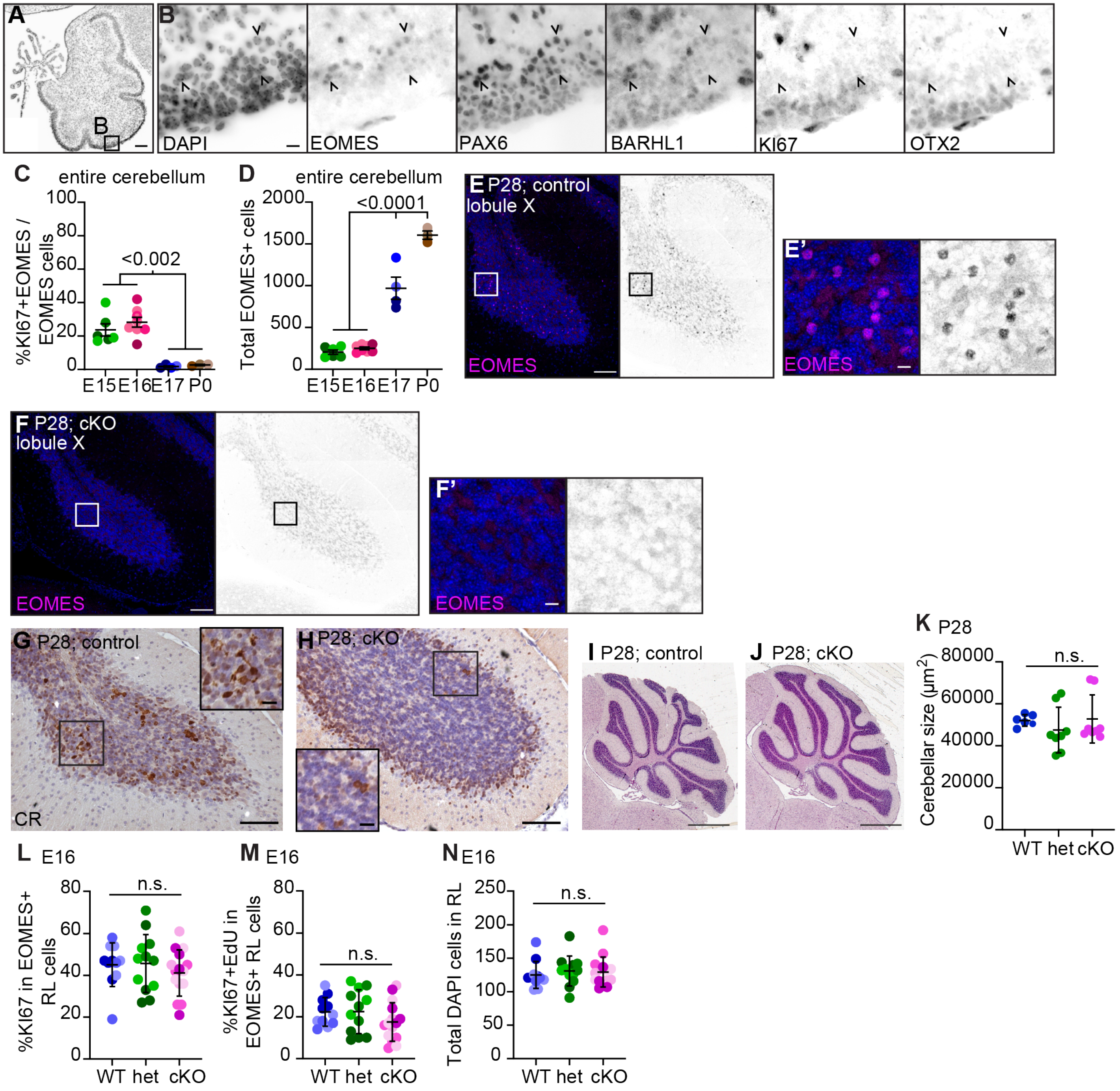


**Figure S2. cKO of *Eomes* DNA-binding domain in the *Lmx1a-Cre* lineage results in lack of mature UBCs.**
**A**) DAPI image of P0 cerebellum. Scale bar, 100 µm. **B**) Zoom of box in A showing Immuno-SABER of indicated antibodes. Carets align EOMES$^{lo}$ post-mitotic granule cells in the developing posterior lobe. Scale bar, 10 µm. **C**) Quantification of Immuno-SABER %KI67+EOMES/EOMES+ cells in cerebellum over time. Ordinary one-way ANOVA. Multiple comparisons revealed E15 and E16 as significantly different from E17 and P0, with displayed p-values. **D**) Quantification of Immuno-SABER total EOMES+ cells in cerebellum over time. Ordinary one-way ANOVA. Besides E15 and E16, all time points were significantly different from each other, $p<0.0001$. **E,F**) IF image of EOMES (magenta) staining in lobule X in E) *Eomes*$^{control}$ and F) *Eomes*$^{cKO}$ P28 animals. Scale bar, 100 µm and 10 µm (zoom). **G,H**) Immunohistochemistry staining of CR in P28 lobule X of G) *Eomes*$^{control}$ and H) *Eomes*$^{cKO}$ cerebellum. Scale bar, 100 µm and 20 µm (zoom inset). **I,J**) H&E staining of P28 I) *Eomes*$^{control}$ and J) *Eomes*$^{cKO}$ cerebellum. Scale bar, 1000 µm. **K**) Quantification of cerebellar size at P28 in displayed genotypes. n.s., non-significant. **L-N**) Quantification of given genotypes at E16 for L) %KI67 in EOMES progenitor cells in the RL, M) %EdU+KI67 in EOMES+ cells in the RL, and N) the total number of RL cells. Ordinary one-way ANOVA. n.s. not significant

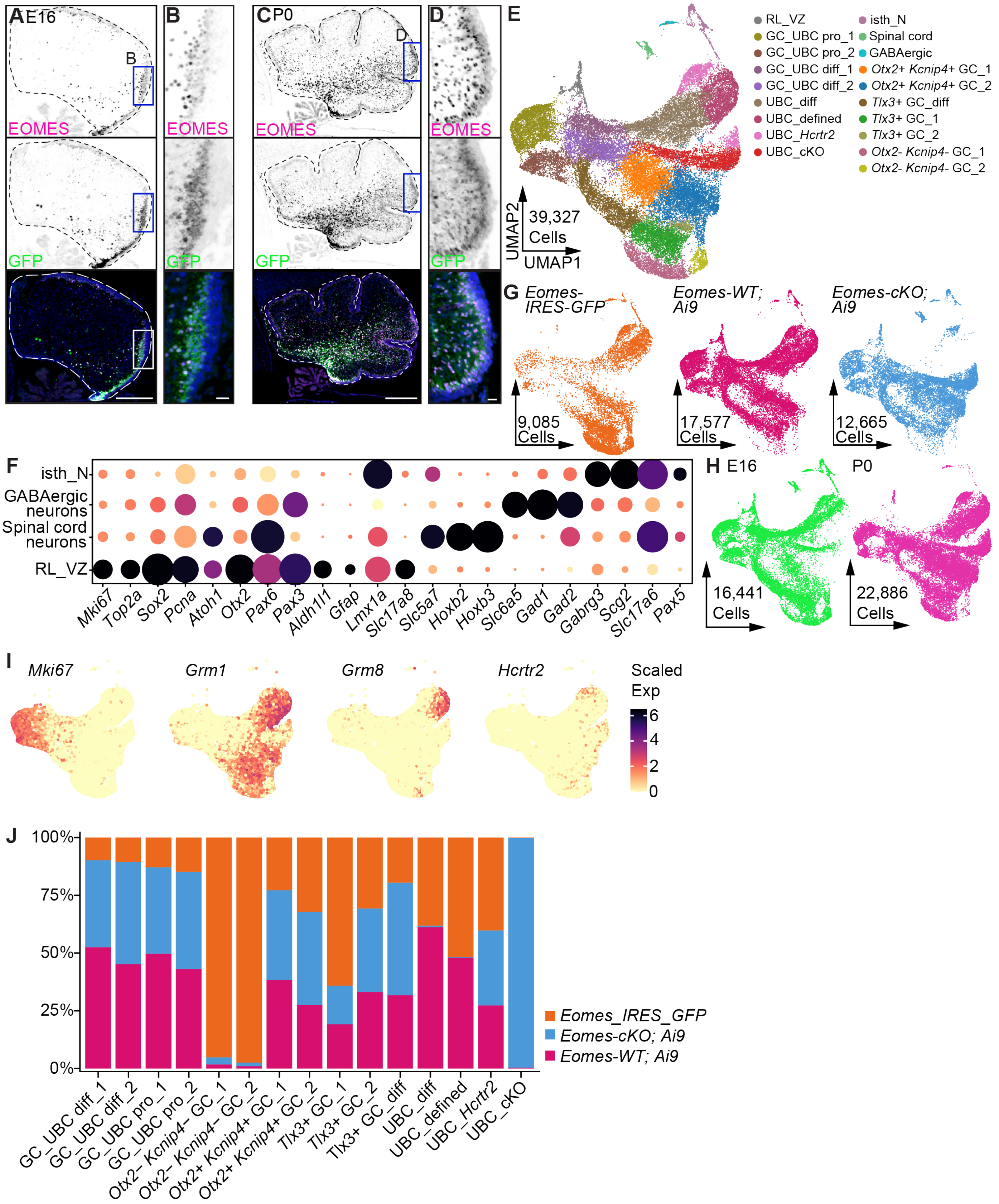


**Supplemental Figure S3** *legend next page* ►

◀*cont. from previous page*

**Figure S3. *Eomes* is expressed in UBCs and developing GCs**. **A**) Representative image of EOMES (magenta) and GFP (green) in *Eomes-IRES-GFP* E16 cerebellum. Scale bar, 200 µm. **B**) Zoom of inset of A. Scale bar, 20 µm. **C**) Representative image of EOMES (magenta) and GFP (green) from *Eomes-IRES-GFP* P0 cerebellum. Scale bar, 200 µm. **D**) Zoom of inset of C. Scale bar, 20 µm. **E**) Cells recovered in single-cell RNA-seq experiment after removal of low-quality clusters and "*undefined*" cluster consisting of 29 cells. RL_VZ, rhombic lip ventricular zone. Isth_N, isthmic neurons. **F**) Marker gene expression of non-GC_UBC clusters, which were removed from further analysis. **G**) Cells contributed by genotype. **H**) Cells contributed by age. **I**) Scaled expression of additional marker genes. **J**) Proportion of cells contributed to each GC_UBC lineage cluster in the displayed genotypes.

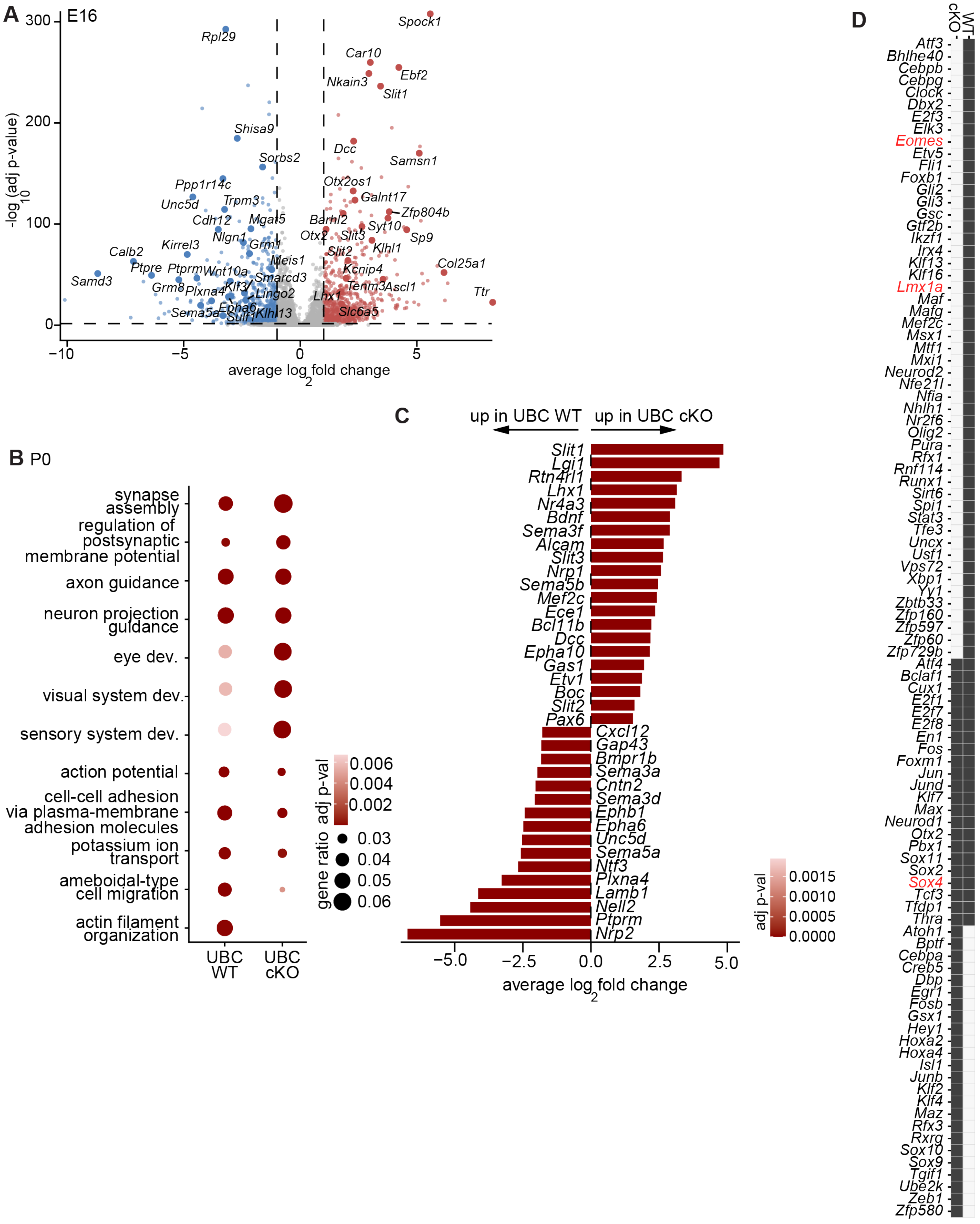


**Supplemental Figure S4** *legend next page* ▶

◀*cont. from previous page*

**Figure S4. *Eomes* is required for UBC maturation and migration.**
**A**) Volcano plot of pseudobulked UBC_cKO vs UBC at E16. Blue, genes downregulated in UBC_cKO. Pink, genes upregulated in UBC_cKO. **B**) Gene ontology analysis of pathways up and downregulated in UBC_cKO at P0. **C**) Specific axon guidance-related genes expressed in wild type UBCs (left) and UBC_cKOs (right). **D**) All SCENIC active gene regulatory networks (black box) present in *Eomes-WT;Ai9* and *Eomes-cKO;Ai9* cells.

**Table S1. Proteins targeted using Immuno-SABER.**

| Target | Cell types |
|---|---|
| EOMES | GC_UBCP, mature UBCs |
| LMX1A | RL progenitors, GC_UBCP, UBCs |
| BARHL1 | Glutamatergic lineage |
| ATOH1 | RL progenitors, EGL/GCPs |
| OTX2 | Posterior GC/Ps, UBCs |
| PAX6 | Glutamatergic lineage |
| SOX2 | Ventricular zone progenitors |
| KI67 | Cell cycle |
| mGluR1$\alpha$ | Type II UBCs |
| Calretinin | Type I UBCs |
| TRPC3 | Type II UBCs |

Because of limited amount of working antibody, ATOH1 and LMX1A were not included in all samples. SOX2, and OTX2 were visualized after Immuno-SABER with traditional antibody staining (secondaries against rat and goat, respectively). See *Methods* for details.

**Table S2.** Differentially expressed genes ($\log_2$ fold change > |1|, adjusted p-value <0.05) between pseudobulked UBC and UBC-cKO clusters at E16 and P0.

**Table S3.** Gene set enrichment analysis of the pseudobulked UBC and UBC-cKO clusters at E16 and P0.

**Table S4.** Gene ontology terms for enriched biological processes for pseudobulked UBC and UBC-cKO clusters at E16 and P0.
